# Hypoxic memory of tumor intrinsic type I interferon suppression promotes breast cancer metastasis

**DOI:** 10.1101/2022.05.12.491632

**Authors:** Oihana Iriondo, Desirea Mecenas, Yilin Li, Christopher R. Chin, Amal Thomas, Yonatan Amzaleg, Aidan Moriarty, Veronica Ortiz, Matthew MacKay, Amber Dickerson, Grace Lee, Sevana Harotoonian, Bérénice A. Benayoun, Andrew Smith, Christopher Mason, Evanthia T. Roussos Torres, Remi Klotz, Min Yu

## Abstract

Hypoxia is a common feature of many solid tumors due to aberrant proliferation and angiogenesis and has been associated with tumor progression and metastasis. Most of the well-known hypoxia effects are mediated through hypoxia-inducible factors (HIFs), but the long-lasting effect of hypoxia beyond the immediate HIF regulation remains less understood. Here we show that hypoxia exerts a prolonged effect to promote metastasis. Using breast cancer patient-derived circulating tumor cell (CTC) lines and common breast cancer cell lines, we found that hypoxia downregulates tumor intrinsic type I interferon (IFN) signaling and its downstream antigen presentation (AP) machinery in luminal breast cancer cells, via both HIF-dependent and HIF-independent mechanisms. Hypoxia induced IFN/AP suppression can last longer than the hypoxic exposure, presenting a “hypoxic memory” phenotype. Hypoxic memory of IFN/AP downregulation is established by specific hypoxic priming, and cells with hypoxic memory have an enhanced ability for tumorigenesis and metastasis. The histone deacetylase inhibitor (HDACi) Entinostat can erase the hypoxic memory and improve the immune clearance of tumor cells when combined with checkpoint immunotherapies in a syngeneic breast cancer mouse model. These results point to a novel mechanism for hypoxia facilitated tumor progression, through a long-lasting memory that provides advantages for CTCs during the metastatic cascade.

**Significance:** We revealed a novel hypoxic memory of prolonged suppression of tumor intrinsic type I IFN and AP signals that promote tumorigenesis and metastasis, suggesting novel mechanistic understanding of the immune evasive properties of CTCs.

## Introduction

In solid tumors, rapid cell division and aberrant angiogenesis often lead to oxygen deprivation or hypoxia, which has long been associated with increased malignancy, poor prognosis, and resistance to radio- and chemotherapy(1). Hypoxia influences tumor progression through a variety of mechanisms (reviewed in(2)), including promotion of angiogenesis(3), matrix remodeling(4), enhanced cell motility and cancer stem cell content(5–7), and modification of the premetastatic niche(8,9). Hypoxia-inducible factors (HIFs) play a central role in many of these processes. HIFs are heterodimers composed of an oxygen labile alpha subunit (HIF1α, HIF2α, or the lesser known HIF3α) and a stable beta subunit (HIF1β, also known as ARNT). Under well oxygenated conditions, HIF1α and HIF2α are hydroxylated in two conserved proline residues by EGLN proteins (also known as prolyl hydroxylases, PHDs), bound by the von Hippel-Lindau (VHL) protein and targeted for proteasomal degradation(10–12). PHDs belong to the 2-oxoglutarate-dependent dioxygenase (2OGDD) superfamily and use O_2_ as a co-substrate.

Therefore, when oxygen availability is low HIFα is stabilized, binds to HIF1β and regulates the expression of a myriad of genes involved in proliferation, metabolism, cell motility, apoptosis, etc.(13,14). Several studies have underscored the importance of alternative HIF-independent mechanisms in regulating the cellular response to hypoxia. Besides HIFα, PHDs can hydroxylate non-HIFα substrates in an oxygen-dependent manner, regulating their stability or activity(15). In addition, other enzymes from the 2OGDD superfamily of proteins have been shown to function as oxygen sensors, including TET DNA demethylases(16) and several Jumonji (JmjC) histone demethylases (KDMs)(17,18). Furthermore, some TET and JmjC-KDMs are transcriptionally regulated by HIF(19), highlighting a complex role for hypoxia in shaping the epigenetic landscape of cells.

Tumor hypoxia has been linked to immunosuppression (20,21). Hypoxic areas in tumors attract M2-like tumor associated macrophages (TAMs)(22,23), myeloid-derived suppressor cells (MDSCs)(24) and regulatory T cells (Tregs), and show reduced CD8 T cell infiltration(25). Furthermore, hypoxic microenvironments can lead to diminished T cell performance and survival, mostly due to metabolic alterations driven by hypoxia, such as increases in lactic acid and extracellular adenosine(20,26). However, the effect of hypoxia on tumor cell intrinsic immune evasive features are less characterized.

A key class of cytokines that play pivotal roles in the immune response against infectious agents and tumors are type I interferons (IFNs)(27). Type I IFNs are a family of protein consisting of 13 IFNα subunits, IFNβ, and several poorly defined single gene products(28). Expression of type I IFNs is induced by the activation of pattern and danger recognition receptors. Once secreted, IFNs signal through the IFNα/ý receptor (IFNAR, composed of subunits IFNAR1 and IFNAR2), which phosphorylates JAK1 and TYK2, resulting in phosphorylation and activation of STAT1/STAT2 heterodimers. Phosphorylated STAT1/STAT2 bind to IRF9 to form the interferon-stimulated gene factor 3 (ISGF3). ISGF3 translocates to the nucleus and binds to interferon signaling response elements (ISRE) to induce the expression of interferon stimulated genes (ISGs). In addition to this canonical signaling, type I IFNs can activate signaling by STAT3-6, JNK, ERK, MAPK and mTOR pathways(28). Type I IFNs can regulate tumor growth directly – by impacting cell proliferation and survival – or indirectly – by modulating antitumor immunity or influencing tumor angiogenesis(29).

Our previous work has shown that the hypoxia signaling pathway is significantly upregulated in CTCs isolated from patient blood during disease progression compared to those isolated during treatment response(30). Hypoxia is a condition that facilitates the establishment of CTC lines(30,31) with the capacity to metastasize to the major organs of experimental mice, as seen in the corresponding patients(32). Furthermore, recent reports showed that hypoxia promotes CTC generation and pro-metastatic behavior(33,34). It has been clear that hypoxia induced molecular changes could play important roles in promoting metastasis. However, the long term effect of hypoxia in CTCs that exited the exposure has not been fully characterized.

## Results

### Hypoxia downregulates IFN and AP pathways in luminal breast cancer cells

To test if hypoxic cultures impact the tumor formation and metastatic capacity of CTC lines, we grew GFP-LUC labeled CTCs for 8 weeks in normoxic or hypoxic cultures, and then injected them into the mammary fat pads or the tail veins of NSG mice (Figure 1A). Once the CTC lines are established, normoxic culture did not decrease cell growth (Figure S1A) nor significantly affect cell death (Figure S1B). *In vivo*, BRx50 and BRx68 CTC lines formed larger tumors after being cultured in hypoxia compared to normoxia, while BRx07, BRx42 and BRx61 cells formed similarly sized tumors following culture in both conditions (Figures 1B and S1C). In the setting of experimental metastasis via tail vein injection, 7 out of 10 mice injected with BRx68 cells grown in hypoxic conditions showed metastatic signals detected by bioluminescent imaging 14 weeks after injection, compared to only 1 out of 7 mice injected with BRx68 cells grown in normoxia (Figures 1C and S1D). In contrast, mice injected with BRx07 cells cultured in normoxia or hypoxia showed similar levels of metastasis (Figure S1E).

**Figure 1:**
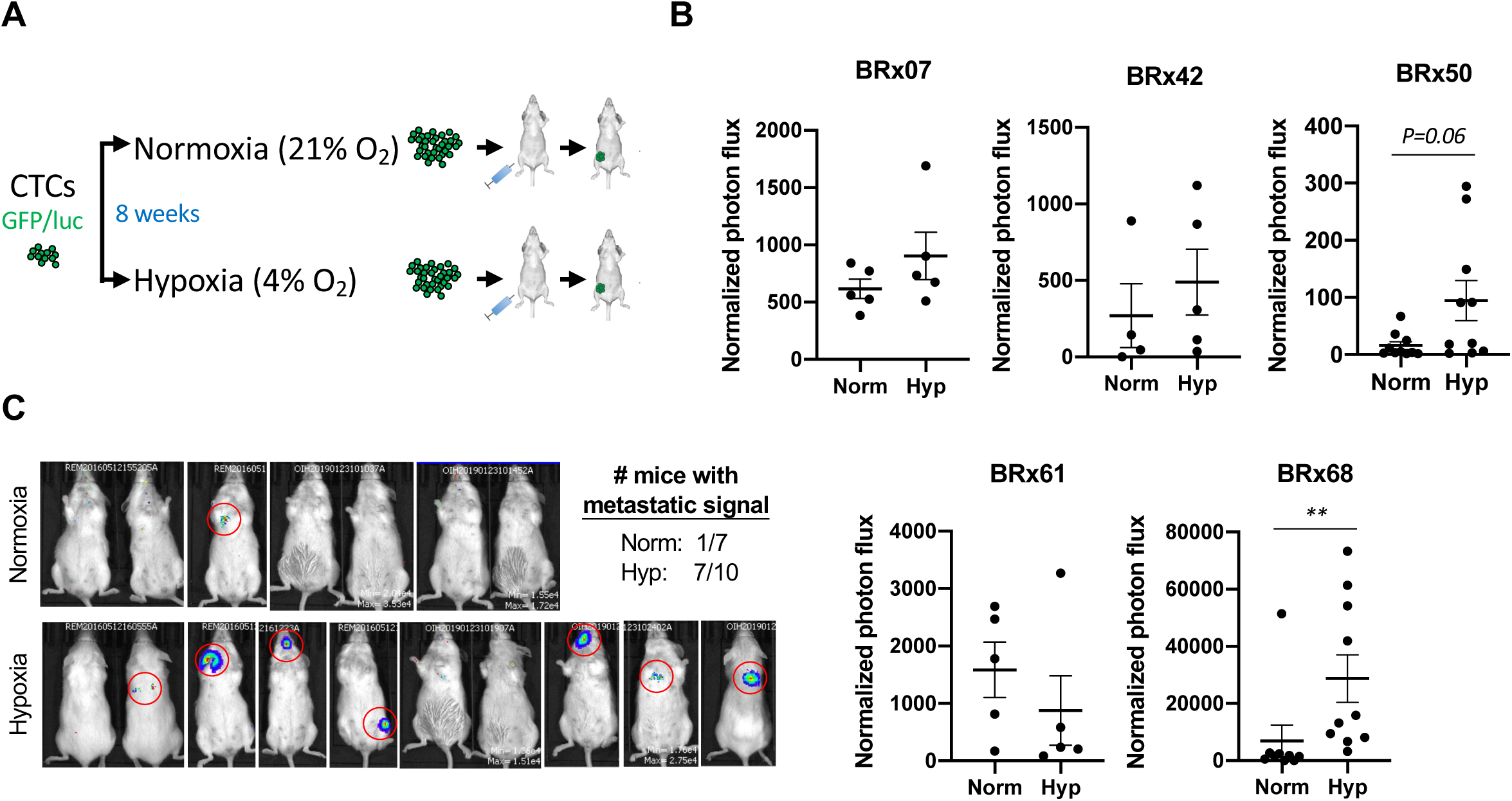
Hypoxia promotes primary tumor growth and metastasis formation in certain breast cancer CTC lines. **A)** Schematic representation of xenotransplantation experiments. Cells constitutively expressing GFP and luciferase (GFP/luc) were grown in normoxia or hypoxia for 8 weeks and injected into the 4^th^ mammary fat pads or tail veins of NSG mice. **(B)** Tumor growth in mice orthotopically injected with different CTC lines (2,000 cells per mouse for BRx07, BRx61 and BRx68, 20,000 cells per mouse for BRx42 and BRx50), measured by bioluminescence. Normalized photon flux at 12 weeks after injection. Means ±SEM are shown. n = number of mice. P values were obtained with 2-tailed Mann-Whitney test. *P<0.05, **P<0.01. (**C)** Bioluminescence imaging data of mice 14 weeks after intravenous injection of 100,000 BRx68-GFP/luc cells per mouse.

RNA-seq analysis of BRx68 cells cultured in normoxia or hypoxia for 8 weeks showed 84 genes upregulated and 241 genes downregulated in hypoxia (fold change≥1.5, FDR≤0.05) (Figure 2A; Table S1-S2). Pathway analysis of these differentially expressed genes showed an enrichment for immune related pathways, especially interferon (IFN) signaling and the antigen presentation (AP) pathway. All genes in these pathways showed lower expression in hypoxia (Figure 2B).

**Figure 2:**
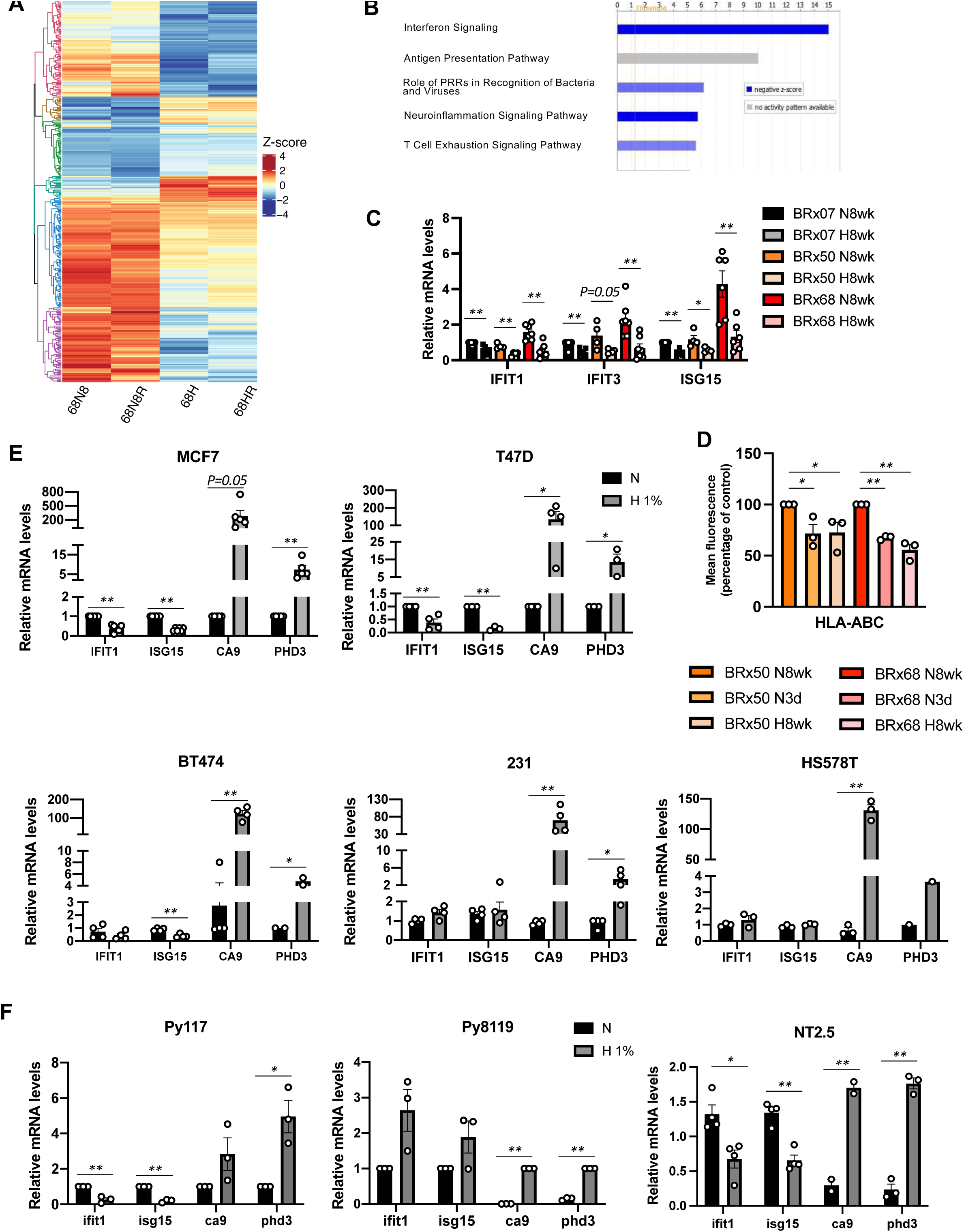
Hypoxia downregulates type I IFN and AP pathways in luminal breast cancer cells. (A) Heatmap depicting genes differentially expressed in BRx68 cells grown in normoxia (N) or hypoxia (H, 4% O_2_) for 8 weeks. **(B)** Top canonical pathways enriched in genes differentially expressed in BRx68 cells cultured in normoxia or hypoxia (4% O_2_). Graph shows -log(p-value). Blue bar indicates negative z-score. **(C)** qPCR analysis of 3 IFN target genes in CTC lines grown in normoxia or hypoxia (4% O_2_) for 8 weeks. Means ±SEM of 7 (BRx07 and BRx68) or 4 (BRx50) experiments are shown. P value was calculated with 2-tailed Student’s *t* test. *P<0.05, **P<0.01. **(D)** Flow cytometry analysis of relative level of HLA-ABC in BRx68 and BRx50 cells cultured for 8 weeks in normoxia (N8), 3 days in normoxia (N3), and 8 weeks in hypoxia (H8). Means ±SEM, n=3, P value was calculated with one-way ANOVA followed by Fisher’s LSD test. *P<0.05, **P<0.01. **(EF)** qPCR analysis of 2 IFN target genes (IFIT1, ISG15) and 2 HIF target genes (CA9, PHD3) in different breast cancer cell lines of human **(E)** or mouse **(F)** origin cultured in normoxia or hypoxia (1% O_2_) for 3 days. Means ±SEM. P values were calculated with 2-tailed Student’s *t* test * p<0.05, ** p<0.01.

Similarly, Gene Set Enrichment Analysis (GSEA) with genes downregulated in hypoxia showed significant overlaps with several gene sets involved in interferon response (Table S3). As expected, the list of genes upregulated in hypoxia significantly overlapped with several hypoxia gene sets (Table S4). qPCR analysis confirmed the lower expression of the IFN target genes ISG15, IFIT1 and IFIT3 in BRx68 cells grown in hypoxia and showed similar results with BRx07 and BRx50 cells (Figure 2C). Flow cytometry analysis of the MHC-I molecules in BRx68 and BRx50 also showed significantly reduced levels in cells cultured in hypoxia, compared to those cultured in normoxia for 8 weeks (Figure 2D).

We next set out to determine whether hypoxia reduces the expression of IFN target genes in other breast cancer cell lines. In response to hypoxia, all human cell lines tested upregulated the HIF target genes CA9 and PHD3 (Figure 2E). However, hypoxia reduced IFIT1 and ISG15 expression in MCF7, T47D and BT474 cells, while failing to do so in MDA-MB-231 and HS578T cells (Figure 2E). We also tested mouse mammary tumor cell lines in hypoxia, and found reduced Ifit1 and Isg15 expression levels in Py117(35) and NT2.5(36) cells, but not in Py8119(35) cells (Figure 2F). To investigate the molecular characteristics distinguishing cells that decrease IFN target gene expression in hypoxia (responders) from those that do not (non-responders), we analyzed publicly available RNA-seq datasets of human breast cancer cell lines and microarray data from mouse cell lines. Genes differentially upregulated in responders showed significant overlap with several sets of genes upregulated in luminal tumors (compared to basal tumors) and genes downregulated during epithelial to mesenchymal transition (EMT) (Tables S5). In contrast, genes overexpressed in non-responders overlapped significantly with sets of genes downregulated in luminal tumors and genes upregulated during EMT (Tables S6).

To evaluate whether the same patterns also presented *in vivo*, we next investigated whether tumors from breast cancer patients showed inverse correlations between a hypoxia signature(37) and a signature composed of genes from the IFN and AP pathways that were differentially expressed in our RNA-seq analysis (IFN/AP signature) (Figure 2A; Table S7). Most breast tumors in the TCGA dataset had low scores for both signatures. However, tumors with high scores for the hypoxia signature generally showed low scores for the IFN/AP signature, and vice-versa (Figures 3A and S2A). Since TCGA datasets are obtained from bulk tumors, the inverse correlation between hypoxia and IFN/AP signatures could be a consequence of the differential recruitment of immune cells to hypoxic vs. non-hypoxic tumors. To explore possible correlations between these 2 signatures in tumor cells *in vivo*, we performed single-cell RNA sequencing (scRNAseq) analysis of spontaneous breast tumors formed in the FVB-MMTV-PyMT mouse model. Using the same mouse model, another group reported that about 2 percent of tumor tissue dissected from 11-weeks-old mice and 10 percent from 16-weeks-old mice were hypoxic(38).

**Figure 3:**
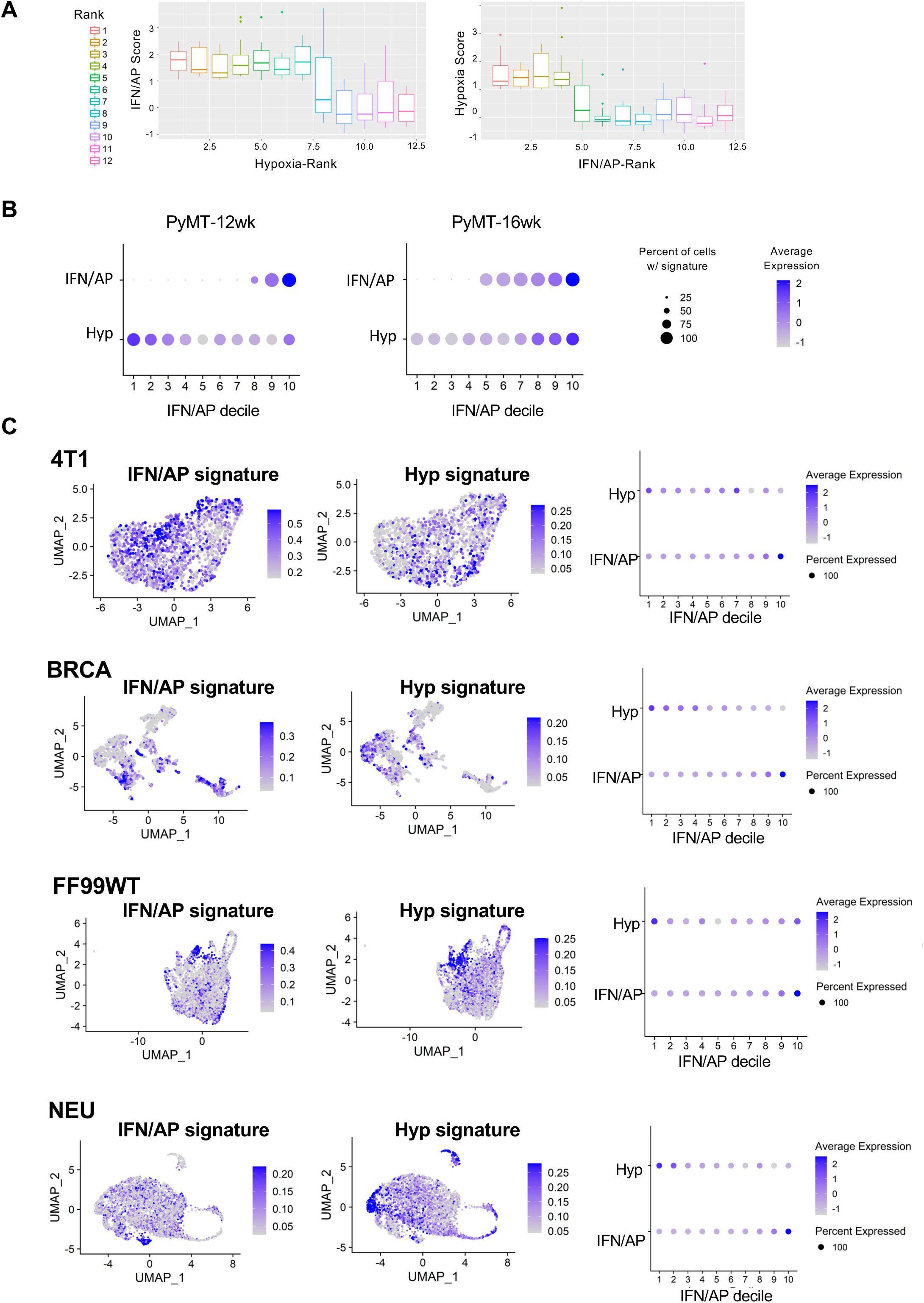
Inverse correlation of hypoxia and IFN/AP signatures in *in vivo* breast tumors and tumor cells of mouse mammary tumors. **(A)** Graph illustrating inverse correlation between hypoxia signature and IFN/AP signature in breast tumors from TCGA. Tumors with detectable scores of hypoxia signature or IFN/AP signature based on RNA-seq analysis were plotted according to the hypoxia score rank (X-axis, Hypoxia-Rank) and IFN/AP Score (Y-axis) or vice versa. **(B)** Decile maps of tumor cell population based on hypoxia and IFN/AP scores for MMTV-PyMT tumors at 12 weeks or 16 weeks. Cell deciles were ranked by IFN/AP scores. **(C)** UMAP plots (left) and decile maps (right) of tumor cell population based on IFN/AP and hypoxia scores for murine breast cancer syngeneic models. Cell deciles were ranked by IFN/AP scores.

Taking this into consideration, tumors from two 12-week-old and two 16-week-old mice were dissociated, and cells were subjected to scRNAseq using the Chromium platform. Using Seurat, we first annotated clusters representing different cell types found in the tumor (Figure S2B), and then projected the hypoxia and IFN/AP signatures on the tumor cells (Figure S2C). For each time point, tumor cells were arranged in deciles according to their IFN/AP scores (Figure 3B). The hypoxia scores for the corresponding tumor cells in each decile showed an inverse correlation with the IFN/AP scores at the earlier time point (12 weeks), but a positive correlation at the later time point (16 weeks) (Figures 3B and S2D). This was consistent with our *in vitro* finding, given reports that the MMTV-PyMT model progresses with a loss of luminal features over time(39). We also applied the same analysis to a published scRNAseq dataset that includes several syngeneic tumor models(40), and found that all 4 models showed a clear inverse correlation of hypoxia and IFN/AP signatures in Cd24-sorted tumor cells (Figure 3C).

### Hypoxic memory of prolonged suppression of IFN/AP is associated with enhanced tumor progression

Intrigued by the differences between BRx07 and BRx68 *in vivo*, we assessed the stability of the hypoxia-induced downregulation of IFN target genes upon reoxygenation. In both BRx07 and BRx68 cells, 3-day reoxygenation results in complete loss of HIF1α expression and decreases HIF target gene expression to same levels as those found after reoxygenation for 7 days (Figures S3A-F). Interestingly, a 3-day reoxygenation period is sufficient to significantly increase IFIT1 and IFIT3 expression levels in BRx07 cells, but not in BRx68 cells (Figures 4A and S3C, S3F). In fact, the increase in IFN target genes observed in BRx07 cells reoxygenated for 3 days was bigger than the increase in cells reoxygenated for 8 weeks (2.9x vs 1.6x for IFIT1, 4x vs 1.75x for IFIT3) (Figure 4A). In BRx68 and BRx50 cells, the increase in IFN target gene expression was only observed when cells were reoxygenated for 8 weeks (Figures 4A and S3G).

**Figure 4:**
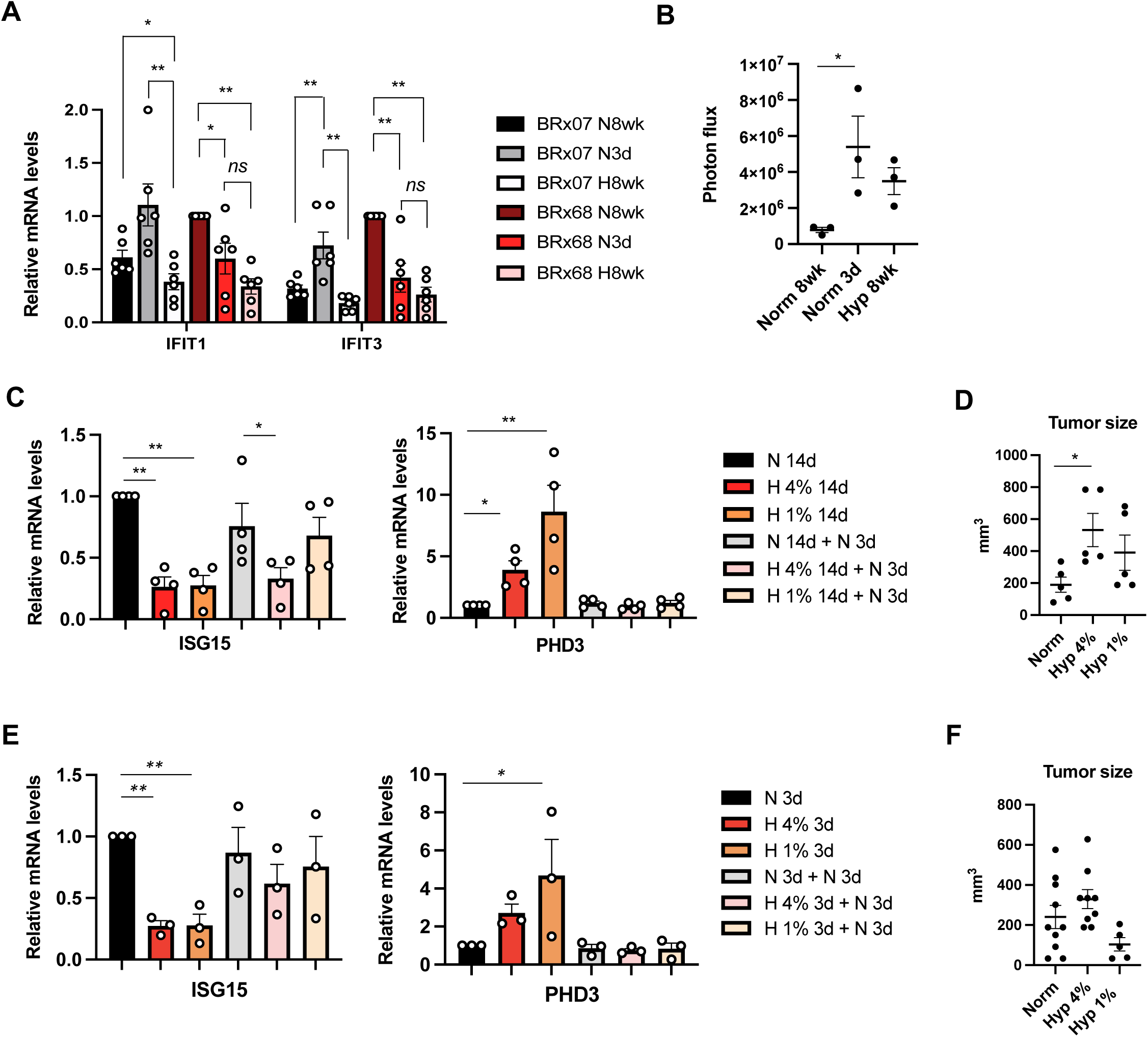
Hypoxic memory of type I IFN suppression is associated with enhanced tumorigenesis and metastasis. **(A)** Relative mRNA expression levels of IFIT1 and IFIT3 in BRx07 and BRx68 cells grown in hypoxia for 8 weeks (4% O_2_, H8), or in normoxia for 8 weeks (N8) or 3 days (N3). Means ±SEM, n=6, P value was obtained with one-way ANOVA followed by Fisher’s LSD test. *P<0.05, ** P<0.01. **(B)** Bioluminescence values from primary tumors 6 weeks after orthotopic injection of 2,000 BRx68-GFP/luc cells. Prior to injection, cells were grown in hypoxia (4% O_2_) for 8 weeks, normoxia for 8 weeks, or normoxia for 3 days. Means ±SEM, n=3. P values were calculated with one-way ANOVA followed by Fisher’s LSD test. *P<0.05. **(C)** qPCR analysis of relative mRNA expression levels of the IFN target ISG15 and the hypoxia target PHD3 in MCF7 cells grown in normoxia or hypoxia (4% O_2_ or 1% O_2_) for 14 days followed by an additional 3 days of reoxygenation. Means ±SEM, n=4. P values were obtained with one-way ANOVA followed by Fisher’s LSD test. *P<0.05, ** P<0.01. **(D)** Tumor volumes formed by MCF7 cells cultured in normoxia or hypoxia (4% or 1%) for 14 days. Means ±SEM, n=5, P values were obtained with one-way ANOVA followed by Fisher’s LSD test. *P<0.05. **(E)** qPCR analysis of relative mRNA expression levels of the IFN target ISG15 and the hypoxia target PHD3 in MCF7 cells grown in normoxia or hypoxia (4% O_2_ or 1% O_2_) for 14 days followed by an additional 3 days of reoxygenation. Means ±SEM, n=4. P values were obtained with one-way ANOVA followed by Fisher’s LSD test. *P<0.05, ** P<0.01. **(F)** Tumor volumes formed by MCF7 cells cultured in normoxia or hypoxia (4% or 1%) for 14 days. Means ±SEM, n=5, P values were obtained with one-way ANOVA followed by Fisher’s LSD test. *P<0.05.

Accordingly, tumors formed by BRx68 cells grown in normoxia for 3 days were significantly bigger than those formed by cells grown in normoxia for 8 weeks (Figure 4B). Similar maintenance of IFN pathway suppression was also observed in HLA-ABC levels in BRx68 and BRx50 after reoxygenation for 3 days (Figures 2D and S3H).

We next evaluated if the sustained suppression of IFN targets by hypoxia observed in CTC lines also occurs in other breast cancer cell lines. When MCF7 cells were cultured in 4% or 1% O_2_ for 14 days and then reoxygenated for 3 days (see Methods), cells that had been grown in 4% O_2_ maintained lower IFN target levels, even though the HIF target PHD3 reached basal expression levels (Figure 4C). Intriguingly, tumor growth in the mammary fat pads of NSG mice showed an association with the prolonged suppression of the IFN target: MCF7 cells grown in hypoxia, especially at 4% oxygen, for 14 days formed bigger tumors than those grown in normoxia (Figure 4D). In contrast, a shorter exposure to hypoxia for 3 days did not maintain the suppression of the IFN target ISG15 (Figure 4E), and MCF7 cells grown in various oxygen conditions for 3 days produced similar sized tumors (Figure 4F). Taken together, these results suggest that certain breast cancer cells can maintain lower IFN signaling even after they exit hypoxic niches, suggesting the existence of a “hypoxic memory”.

### Both HIF-dependent and -independent mechanisms regulate IFN

To clarify the mechanism by which hypoxia reduces IFN target gene expression in luminal breast cancer cells, we first evaluated whether there are different levels of mitochondrial double strand RNA (dsRNA), an endogenous IFN stimulus, as recently reported(41). Immunofluorescence staining using the J2 antibody, which is commonly used to detect mitochondrial dsRNA, did not detect a significant decrease of signal in MCF7 cells cultured in in 4% or 1% hypoxia for 3 days compared to that in normoxia (Figure S4A). To test whether there were different responses to the same stimulus, we transfected the MCF7 cells that have been cultured in different oxygen levels for 14 days with the viral double stranded RNA analogue poly(I:C) (42) and grew them in the normoxic condition for 6 additional hours before harvesting. We found that indeed the activation of phospho-STAT1 in response to the poly(I:C) stimulus is dampened in cells cultured in hypoxia (Figure S4B).

We then evaluated whether HIF activity is needed and used CRISPR-Cas9 to delete ARNT, the HIF1α and HIF2α heterodimerization partner that is essential for the transcriptional activity of HIFs(43). ARNT knockout (KO) in MCF7 cells led to the inability of hypoxia to induce the expression of NDRG1, a target of both HIF1 and HIF2(44) (Figure 5A-B). However, IFN target gene expression was reduced in response to hypoxia both in control and ARNT-KO cells (Figures 5B and S4C), suggesting that the effect of hypoxia on IFN signaling can be observed independent of HIF. Since there are reported PHD dependent but HIF-independent mechanism of hypoxia influence (45), we next treated MCF7 cells with the PHD inhibitor FG4592(46) that led to HIF1α stabilization and induction of HIF target genes in MCF7 (Figure S4D-E) and T47D cells (Figure S4F-H), showing that the drug is efficient in inhibiting PHD activity. Cells treated with this small molecule also exhibited lower expression of IFN target genes (Figures 5C and S4G). Interestingly, in MCF7 ARNT-KO cells, treatment with FG4592 and DMOG, a 2-OG analogue that leads to PHD inactivation and HIF1α stabilization (47) as unable to reduce IFN target gene expression (Figure 5D), suggesting that the effect that inhibition of PHDs has on IFN signaling is mediated by HIF. We then overexpressed mutated forms of HIF1α and HIF2α, with mutations on two prolyl sites, that render them insensitive to regulation by PHDs in normoxia (48), resulting in HIF1α and HIF2α stabilization (Figure S4I-J) and increased expression of downstream targets (Figure 5E-F) in normoxia conditions . Intriguingly, while mutant HIF1α overexpression suppressed the expression of IFN target genes (Figure 5E) mutant HIF2a overexpression had the opposite effect (Figure 5E-F). These data indicate the existence of both HIF1-dependent and HIF-independent mechanisms for downregulating IFN signaling.

**Figure 5:**
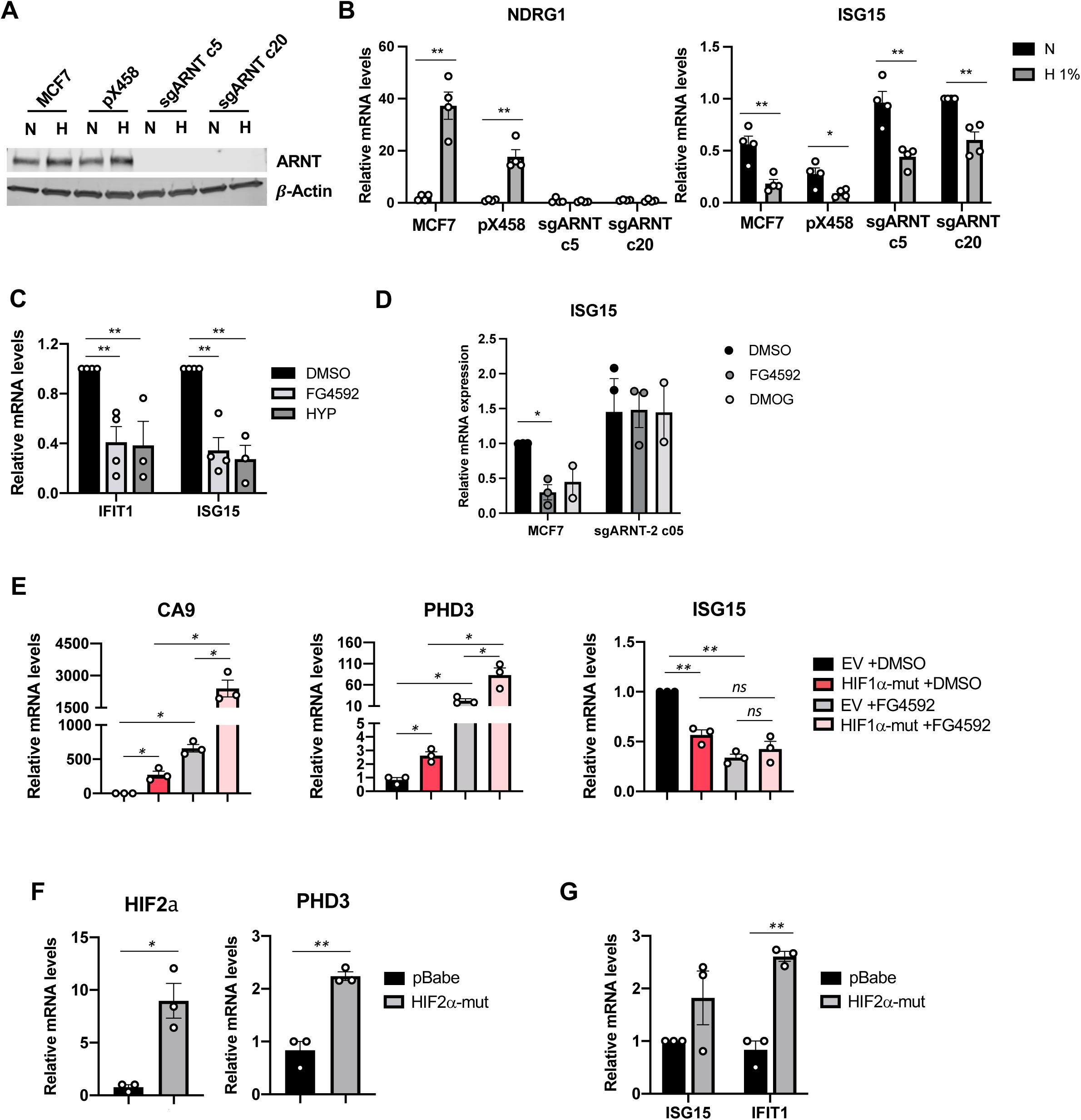
HIF-dependent and -independent mechanisms regulate IFN suppression. **(A)** Immunoblot analysis of ARNT expression in ARNT-KO MCF7 cells, bulk MCF7 and a px458 control MCF7 cells. β-ACTIN is shown as a loading control. **(B)** Relative mRNA expression levels of NDRG1 (left) and ISG15 (right) in bulk MCF7 cells, 1 px458 control MCF7 clone and 2 ARNT-KO MCF7 clones grown in normoxia or hypoxia (1% O_2_) for 3 days. Means ±SEM, n=4. P values were obtained with one-way ANOVA followed by Fisher’s LSD test. **(C)** IFIT1 and ISG15 expression levels in MCF7 cells treated with the PHD inhibitor FG4592 (100 μM) or hypoxia for 3 days. Means ±SEM, n=4. P values were obtained with one-way ANOVA followed by Fisher’s LSD test. *P<0.05, ** P<0.01. **(D)** ISG15 expression levels in MCF7 cells, or MCF7 ARNT-ko cells treated with the PHD inhibitor FG4592 (100 μM) or DMOG for 3 days. Means ±SEM, n=3 for FG4592, n=2 for DMOG. P values were obtained with one-way ANOVA followed by Fisher’s LSD test. *P<0.05. **(E)** Relative mRNA expression levels of CA9 (left), PHD3 (middle) and ISG15 (right) in T47 cells with EV, HIF1α-mutant expression, treated with DMSO or FG4592 in normoxia culture for 3 days. Means ±SEM, n=3. P values were obtained with one-way ANOVA followed by Fisher’s LSD test. *P<0.05, **P<0.01. **(F)** Relative mRNA expression levels of HIF2α (left), PHD3 (middle) and ISGs (right) in MCF7 cells with pBabe-EV or HIF2α-mutant expression in normoxia culture for 3 days. Means ±SEM, n=3. P values were obtained with one-way ANOVA followed by Fisher’s LSD test. *P<0.05, **P<0.01.

We next investigated potential HIF-independent mechanisms for hypoxic suppression of IFN signaling. Recent reports have shown that, similar to the PHDs that hydroxylate HIF in well oxygenated environments, other proteins of the 2-oxoglutarate-dependent dioxygenase family, including some histone demethylases, show low oxygen affinity (high K_M_), enabling them to sense physiological changes in oxygen(18,49). As a result, hypoxic culture decreases the activity of such histone demethylases, leading to increased levels of methylation in several H3 residues (Figure S5A). KDM6A, KDM5A and KDM4B are 3 of the histone demethylases with the highest K_M_ for oxygen(18). To determine if inhibition of these histone demethylases can mimic the effect of hypoxia on interferon target genes, we treated MCF7 cells with small molecule inhibitors that specifically target these enzymes. Treatment with the KDM6 inhibitor GSK-J4(50) increased H3K27 trimethylation, but actually enhanced IFN target gene expression (Figures S5B-C). Similarly, MCF7 cells treated with the KDM5 inhibitor KDM5-C70(51) that increases H3K4me3 showed small but significant increases in IFN target gene expression (Figure S5D-E). The KDM4 inhibitor NSC636819(52) promoted a modest increase in H3K9me3 levels (Figure S5F). NSC636819 treatment failed to mimic the effect of hypoxia on IFN target genes (Figure S5G), although it did downregulate ESR1 and its target gene PS2 (Figure S5H), as reported elsewhere(53,54). These data suggest that the reduced activities of KDMs are not directly responsible for suppressing the IFN/AP signal.

### Hypoxic tracing plasmids allows validation of hypoxic memory *in vivo*

To elucidate whether such “hypoxic memory” can be found *in vivo*, we designed a 2-plasmid reporter system that enables us to permanently mark cells exposed to hypoxia so that they can be tracked during tumor progression (Figure 6A). Cells that express this 2-plasmid reporter system are initially RFP positive but upon exposure to hypoxia, permanently lose RFP expression due to gene excision and start expressing GFP. FACS analysis of MCF7 cells transduced with the 2-plasmid reporter showed that, while the vast majority of cells remained RFP^+^GFP^-^ when cultured in normoxic conditions, an average of 56.7% of cells grown in hypoxia (1% O_2_) for 3 days were GFP^+^ (Figure S6A-B). When the hypoxia tracing MCF7 (MCF7-HypT) cells were cultured in 3D mixed matrigel/collagen gels, spheres that were originally exclusively RFP^+^ developed GFP^+^ areas as they grew, suggesting the system can detect local hypoxia in spheres (Figure S6C). We next injected MCF7-HypT cells into the mammary fat pads of NSG mice. After 8 weeks, mice were injected with the hypoxia detecting probe pimonidazole(55), and tumors and lungs were dissected and fixed 90 minutes later. Immunostaining of tissue sections suggested that cells that have been exposed to hypoxia were more efficient in metastasizing to the lung, since the proportion of GFP^+^ metastatic loci in lungs was significantly higher than the proportion of GFP^+^ cells in primary tumors (Figures 6B and S6D-E). Importantly, immunofluorescence staining also revealed that CRE expression was exclusively detected in pimonidazole-positive areas, while GFP^+^ cells could be detected in non-hypoxic areas, showing that many cells had exited hypoxic niches after converting into GFP-expressing cells (Figure 6C. In fact, staining of whole tumor sections demonstrated that the vast majority of GFP^+^ cells were located in pimonidazole-negative non-hypoxic regions of the tumor (Figure S6F). In accordance with our *in vitro* results, RFP^-^ GFP^+^ cells sorted from MCF7-HypT tumors showed lower expression levels of IFN target genes than their RFP^+^GFP^-^ counterparts, even though the expression of the HIF target genes CA9, PHD3 and NDRG1 was not significantly different (Figures 6D and S6G), suggesting that cells retain a memory of their hypoxic past.

**Figure 6:**
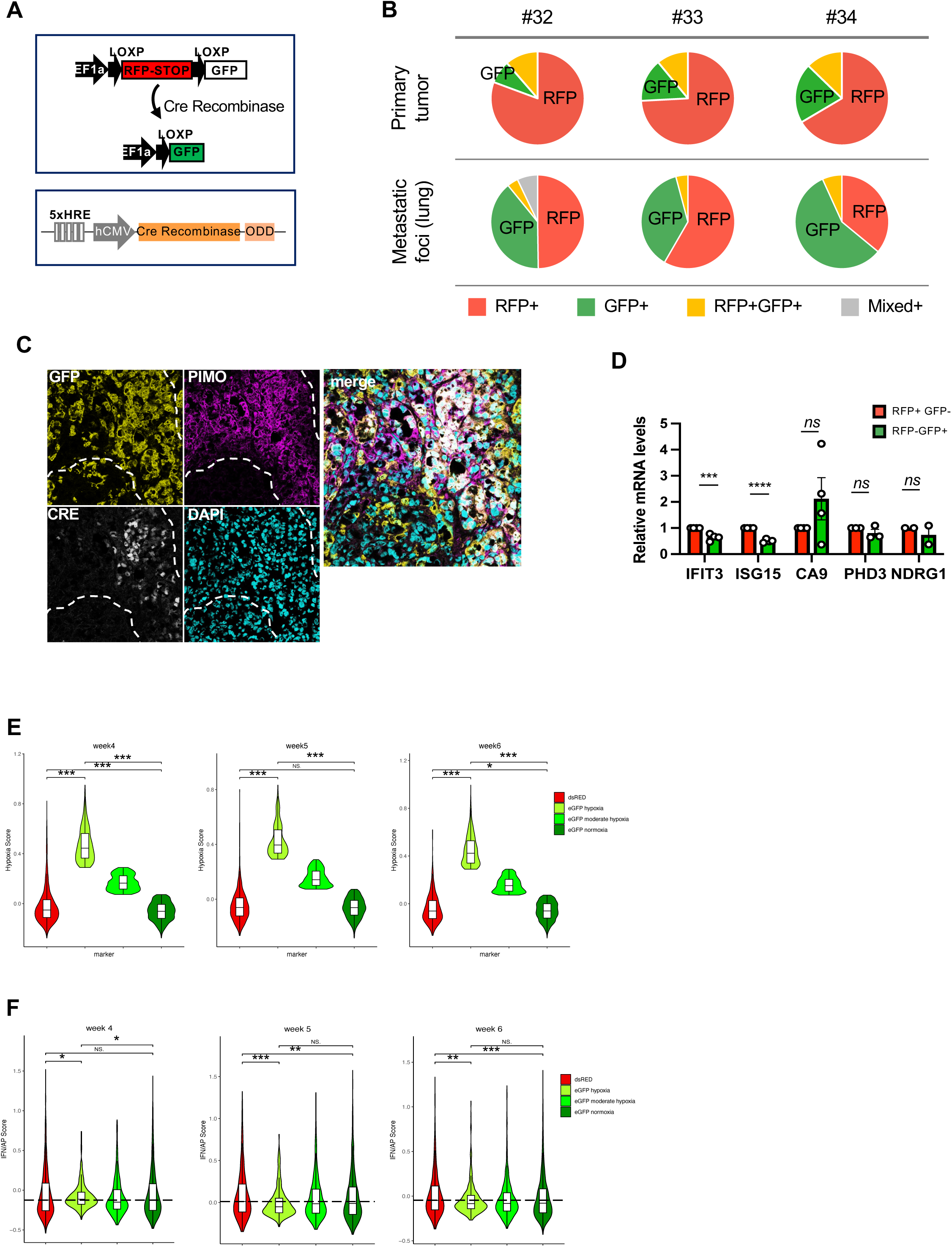
Hypoxia-tracing reporter system showed tumor cells with hypoxic memory and enhanced metastatic capacity. **(A)** Schematic representation of the plasmids used to create the hypoxia tracing MCF7-HypT cells. **(B)** Quantification of RFP and GFP positive area in primary tumors (top) and metastatic loci in lungs (bottom). Mouse number is labeled on the top. **(C)** Representative fluorescent images of a MCF7-HypT tumor section stained with antibodies against GFP (green), pimonidazole (white) and Cre (red). Nuclei were stained with DAPI (blue). Dotted line contours the hypoxic region. Scale bars = 100 μm. **(D)** Relative mRNA levels of IFN targets (IFIT3 and ISG15) and hypoxia targets (CA9, PHD3 and NDRG1) in RFP+GFP-and RFP-GFP+ cell populations sorted from MCF7-HypT tumors. Means ±SEM, n=4, P values were obtained with 2-tailed Student’s t test. *P<0.05, ns: no significance. **(E-F)** Violin plots of hypoxia score **(E)** and IFN/AP score **(F)** from the scRNAseq data in RFP (dsRED), eGFP-hypoxia, eGFP-moderate hypoxia, eGFP-normoxia cells at week 4, 5 and 6. *P<0.05, **P<0.01, ***P<0.001. NS: not significance.

In order to evaluate the hypoxic memory *in vivo* at the single cell level, we optimized our HypT plasmid to include a 3’ UTR for the RFP transcript (Figure S6H), in addition to the 3’UTR of the GFP side, and used the 3’capture methods of chromium to analyze the scRNAseq data of MCF7-HypT tumors at 4, 5, and 6 weeks. Tumor cells with detected RFP or GFP transcript were first ranked based on the hypoxia signature score (Figure S6I). Since GFP positive cells could be tumor cells that are in hypoxic region, or have already exited hypoxic region, the hypoxic score could help distinguish the two situations. We termed the “normoxia” population based on the hypoxia scores of zero or below, “hypoxia” population for the highest decile of the hypoxia score, and “moderate hypoxia” for those in between. As expected, majority of the RFP positive cells (dsRed) are in normoxia; GFP-hypoxia indicates cells in hypoxia, and GFP-normoxia indicates cells that exited the hypoxia region (Figure 6E). The IFN/AP scores were analyzed in the corresponding cell populations (Figure 6F). In all 3 time points, we observed statistically significant decreased IFN/AP score in GFP-hypoxia cells compared to dsRed cells. Intriguingly, from week 5 to week 6, we also observed that such decrease is maintained in the GFP-normoxia cells. Therefore, these *in vivo* data supported our *in vitro* finding of hypoxic memory of IFN/AP suppression.

### Erasing hypoxic memory enhances immune clearance when combined with immune checkpoint inhibitors

To investigate the potential involvement of epigenetic regulation in the hypoxic memory of IFN/AP suppression, we evaluated the ability of several epigenetic inhibitors to rescue IFN/AP suppression after reoxygenation. An HDACi, MS275 (Entinostat)(56), and a low concentration of a G9a/GLP inhibitor, UNC0638(57), reversed the hypoxic memory of IFN target IFIT1 (Figures 7A-B and S7A). To evaluate the effect *in vivo*, we performed scRNAseq analysis of the NT2.5 cells in syngeneic tumors treated with vehicle, Entinostat alone, or in combination with anti-CTLA4, anti-PD1, or both(58). A total of 20 tumors (4 from each group) were processed by 10x Chromium platform after 3 weeks of treatment, but before a significant tumor shrinkage. Surprisingly, different from other luminal mouse single cell data (Figure 3B-C), decile plots of the tumor cells ranked by IFN/AP score in the vehicle condition showed a positive correlation of hypoxia and IFN/AP signatures, but Entinostat treatment led to a clear inverse correlation, and the effect became even more obvious when combined with the immunotherapies (Figure 7C). To investigate whether the existence of hypoxic memory of NT2.5 cells could potentially explain this result, we performed analysis of NT2.5 cells for the long term effect of hypoxia. Indeed, NT2.5 cells showed hypoxic memory of IFN target suppression after reoxygenation from both 4% or 1% of hypoxia exposure for 14 days (Figure 7D). It is tempted to speculate that Entinostat removed the hypoxic memory and upregulated the IFN/AP signals for tumor cells in the normoxic region (negative for hypoxia), leading to a clear inverse correlation of hypoxia-IFN/AP scores in the scRNAseq analysis. We also found that the combination of Entinostat with checkpoint blockades led to an overall increased hypoxia score in the tumors (Figure S7B), suggesting more effective elimination of cells in normoxia. We further investigated the hypoxic memory association with tumor growth in a syngeneic model. When NT2.5 cells grown in normoxia or hypoxia (4% or 1% O_2_) for 14 days were injected in immunocompetent FVB mice, cells pre-cultured in 1% hypoxia generated significantly bigger tumors, compared to those pre-cultured in normoxia (Figure 7E), accompanied with more immunosuppressive Arginase (Arg)+ granulocytic myeloid derived suppressor cells (gMDSCs) and macrophages (Figures 7F-G and S7C-F). These data suggest that besides the intrinsic effect on the tumor cells, this hypoxic memory can modify the recruitment of immune cells to shift the balance towards a more immunosuppressive microenvironment.

**Figure 7:**
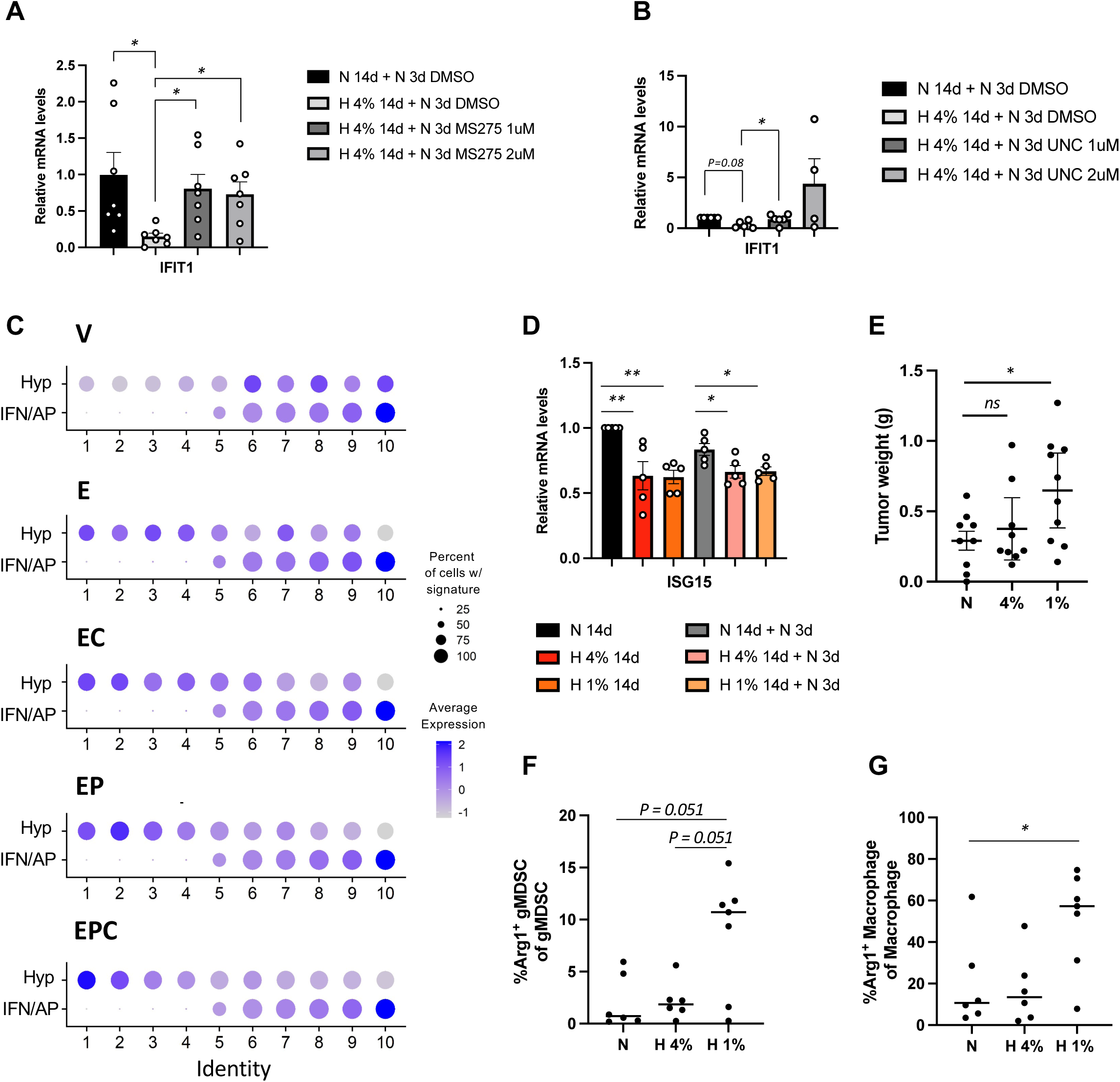
An HDAC inhibitor can erase hypoxic memory and enhance immune clearance of tumor cells in normoxic regions. **(A-B)** Relative mRNA expression levels of IFIT1 in MCF7 cells in reoxygenation experiment (primed in normoxia or hypoxia for 14 days and then reoxygenated in normoxia for 3 days) with or without MS275 **(A)** or UNC0638 **(B)**. n=7, mean±SEM, statistical significance was calculated with Brown-Forsythe and Welch’s ANOVA test followed by unpaired t-test with Welch’s correction, *p<0.05. **(C)** scRNAseq analysis of hypoxia and IFN/AP scores in tumor cells in the syngeneic mammary tumors formed by NT2.5 cells, treated with vehicle (V), Entinostat (E), Entinostat combined with anti-CTLA4 antibody (EC), with anti-PD1 antibody (EP), or with both (EPC). Decile plot of hypoxia and IFN/AP scores for cell population ranked based on IFN/AP scores. **(D)** Relative mRNA expression levels of isg15 in NT2.5 cells cultured in normoxia, 4% or 1% hypoxia for 14 days, and reoxygenated for 3 days. Means ±SEM, n=5, Statistical significance was calculated with one-way ANOVA followed by Fisher’s LSD test. *P<0.05, ** P<0.01. **(E)** Graph showing weight of the orthotopic tumors formed by NT2.5 cells injected into FVB mice at 8 weeks. Means ±SEM. n=9 (Norm), n=9 (Hyp 4%) and n=10 (Hyp 1%). Statistical significance was calculated with one-way ANOVA followed by Fisher’s LSD test. *P<0.05. **(F-G)** Graph showing percentage of Arg+ gMDSCs **(F)** and Arg+ macrophages **(G)** in the same tumors formed by NT2.5 cells in (**E)**. Statistical significance was calculated with nonparametric Mann-Whitney U test. *P<0.05. **(H)** Graph illustrating a proposed model of hypoxia mediated suppression of IFN/AP and the hypoxic memory.

## Discussion

In this study, we discovered that hypoxia leads to the suppression of type I IFN signaling and downstream AP pathway specifically in the luminal subtype of breast cancer cells. This effect is mediated via both HIF1-dependent and HIF-independent mechanisms. Intriguingly, even after a returning to normoxic conditions after a prolonged exposure to low oxygen levels, hypoxia induced IFN/AP suppression can be maintained as a hypoxic memory, which is associated with enhanced tumorigenesis and metastasis. This suggests a novel mechanism for the enhanced metastatic capacity of tumor cells shed from hypoxic regions.

As one of the common tumor microenvironment (TME) features of solid tumors, hypoxia has well-known multifaceted effects that promote tumor progression(2). Our finding on its effect in suppressing tumor cell intrinsic type I IFN and AP signals provides an additional novel mechanism. Type I IFN signaling has intrinsic tumor suppressive effect, by inhibiting proliferation and promoting apoptosis, and it also plays a key role in immune surveillances against tumors(27). Therefore, by downregulating intrinsic type I IFN as well as downstream AP machinery, hypoxia provides tumor cell intrinsic and extrinsic benefits during tumor progression. Of note, a study with similar topic has recently been reported(59), however with different results. Whereas this report showed similar effects in all subtypes(59), we found that this hypoxia mediated suppression of IFN/AP to be specific in luminal subtypes of breast cancer cells with no obvious difference in basal subtype cell lines. The differences may be due to the level of hypoxia used in the experiments. We mainly focused on relatively mild hypoxia condition (4%-1% O_2_) since this is likely to be more common across tumor types(60), while a much lower oxygen levels (1%-0.1%) have been used in the other report(59). Intriguingly, we did observe differences in the long term effect by different hypoxia priming, highlighting the importance in considering the specific niche effects. Moreover, our scRNAseq analysis of the MMTV-PyMT tumor model further supports such effect being more prominent in luminal characteristic tumor cells based on the report that MMTV-PyMT model progresses with a loss of luminal features over time(39). scRNAseq analysis of other syngeneic breast cancer models also showed a clear inverse correlation of hypoxia and IFN/AP scores in tumor cells sorted based on Cd24 surface marker likely enriching luminal type of tumor cells(40). However, the exact mechanisms underlying the luminal versus basal subtypes in terms of their hypoxic responses are not yet clear and warrants further investigation, which may contribute to a better understanding of why luminal tumors are immune “colder” tumors than the basal subtypes.

We also found that hypoxia downregulates IFN/AP signal in both HIF-dependent and - independent manners. An intriguing novel finding is the distinct effect of HIF1α and HIF2α in regulating ISG expression, with HIF1α suppressing whereas HIF2α inducing ISG genes, which may add another layer of heterogeneity pending on the level of HIF1α and HIF2α in a particular cell. Although we have attempted to elucidate the HIF-independent mechanism, the exact regulation is unclear and worth of further investigation. Different from previously mentioned mechanism in which mitochondrial dsRNA quantity is reduced in hypoxia compared to normoxia(41), we did not detect significant differences in the mitochondrial dsRNA levels. Epigenetic factors may play an indirect role since we observed changes in global histone methylations in hypoxia. Epigenetic mechanisms indeed contribute to hypoxic memory maintenances.

Our intriguing finding of hypoxic memory of IFN/AP suppression is, to the best of our knowledge, the first reported long term suppression of such signal. During our manuscript preparation, a study has utilized a similar 2-plasmid Cre-loxP mediated hypoxia tracing system to evaluate the long-term effect of hypoxia *in vivo*(34). The study noted a long-term effect of hypoxia that promotes metastasis, but only utilized the basal subtype of breast cancer cells. In that context, the hypoxic memory mainly persists via upregulated genes that promote cell survival in the presence of high levels of ROS. As we have shown that hypoxia-mediated IFN/AP suppression is prominent in luminal subtype, it is likely that the signals lasting longer as hypoxic memory is subtype dependent and mechanisms could be heterogeneous. A surprising finding is that mild (4% O_2_) and long term (14 days) of hypoxia exposure, but not severe (1% O_2_), nor a short exposure (3 days), lead to the prolonged memory of IFN/AP suppression in MCF7 cells. While in NT2.5 cells, hypoxic memory can be established by both mild and severe hypoxia conditions. The unique hypoxia priming condition suggests specific molecular mechanisms are involved in establishing and/or maintaining such cellular memory in a cell type dependent manner, which warrants further analysis.

Within solid tumors, hypoxic features constantly change due to tumor growth and cellular movement. Hypoxic memory of IFN/AP suppression could contribute to immune evasion and reduce efficacy of checkpoint immunotherapies. As shown in our *in vitro* MCF7 data and *in vivo* data of NT2.5 syngeneic tumors, Entinostat upregulates IFN/AP signals of cells in normoxic regions, resulting in increased hypoxia scores when combined with immunocheckpoint treatment. Despite the limitation of one static time point and the correlative nature, the data suggests the possibility that for NT2.5 cells with hypoxic memory (ie. low IFN/AP score in the normoxia region) in vehicle condition, such memory can be “erased” by Entinostat treatment thereby leading to improved immune clearance of cells in normoxic area by the immune checkpoint therapies. A more direct link with multiple time points will be worth evaluating in future studies. Moreover, our data of the syngeneic model showed that NT2.5 cells pre-conditioned in hypoxia lead to bigger tumor formation, with recruitment of more immunosuppressive cell types in the TME. Furthermore, the prolonged suppression of IFN/AP would likely also contribute to the immune evasive feature commonly seen in CTCs. Indeed, after exposure to hypoxia, only those CTCs with the capacity for a hypoxic memory of IFN/AP suppression showed enhanced tumorigenesis and metastasis. Our study suggests a novel concept of hypoxia induced long term cellular memory as a potential mechanism for promoting tumor progression.

## Additional information

Financial support: The National Institutes of Health (NIH) grant DP2 CA206653 (M.Y.), the Stop Cancer Foundation (M.Y.), the PEW Charitable Trusts and the Alexander & Margaret Stewart Trust (M.Y.), the Department of Defense Era of Hope Scholar Award (M.Y.), the Richard Merkin Foundation (M.Y.), the Broad Fellowship (O.I.), and the NIH grant T90DE021982 (trainees D.M. and Y.A.). The project described was supported in part (core facilities) by award number P30CA014089 from the National Cancer Institute.

Disclosure of potential conflicts of interest: M. Yu. is the founder and director of CanTraCer Biosciences Inc.

## Acknowledgements

We thank the USC Flow Cytometry Facility and USC Molecular Genomics Core for their technical support; D. Ruble and members of CHLA Molecular Pathology Genomics Core for technical assistance; Drs. J. Lang and T. Aguilera for providing cell lines; Drs. Q. Zhang and H. Harada for providing reagents; C. Lytal for editing the manuscript. This research was supported by the following: The National Institutes of Health (NIH) grant DP2 CA206653 (M.Y.), the Stop Cancer Foundation (M.Y.), the PEW Charitable Trusts and the Alexander & Margaret Stewart Trust (M.Y.), the Department of Defense Era of Hope Scholar Award (M.Y.), the Richard Merkin Foundation (M.Y.), the Broad Fellowship (O.I.), and the NIH grant T90DE021982 (trainees D.M. and Y.A.). The project described was supported in part (core facilities) by award number P30CA014089 from the National Cancer Institute. This research was also supported by funds through the Maryland Department of Heath’s Cigarette Restitution Fund Program (CH-649-CRF).

## Author Contributions

O.I. and M.Y. conceived the study. O.I., D.M., Y.L. and M.Y. designed experiments. O.I., D.M., Y.L., A.D., G.L., S.H. and M.Y. performed experiments. O.I., D.M., Y.L., C.R.C., A.T., Y.A., V.O., M.M., A.D., G.L., B.A.B., E.R.T. and M.Y. contributed to analysis and interpretation of data. C.R.C., A.T., Y.A., M.M., A.S. and C.M. conducted computational analyses. E.R.T. provided scRNAseq data of syngeneic tumors. O.I. and M.Y. wrote the manuscript. All authors edited and commented on the manuscript. M.Y. supervised the study.

## Methods

### Cell culture and reagents

CTC lines derived from patients with luminal breast tumors(31) were cultured in ultra-low attachment plates (Corning) with RPMI 1640 supplemented with 20 ng/ml EGF, 20 ng/ml bFGF, 1X B27 and 1X antibiotic/antimycotic. CTCs were maintained in a Nuaire NU-5831 incubator at 4% O_2_ and 5% CO_2_, and media was changed every 3 or 4 days. For normoxia vs hypoxia experiments, equal number of cells were plated and kept in normoxia or hypoxia (4% O_2_) conditions for the indicated duration.

MCF7, MDA-MB-231, HS578T and HEK293T cells were purchased from the American Type Culture Collection and were maintained in DMEM supplemented with 10% FBS. T47D and BT474 cells were provided by Dr. Julie Lang and were cultured in RPMI media with 10% FBS. Py117 and PY8119 cells were provided by Dr. Todd Aguilera and maintained in F12 media containing 5% fetal clone II (Hyclone) and MITO+ (1:1000 dilution, Corning). NT2.5 cells were provided by Dr. Evanthia Roussos Torres and cultured in RPMI media supplemented with 20% FBS, 1.2% Hepes, 1% L-Glutamine, 1% non-essential aminoacids, 1% sodium pyruvate and 0.2% insulin. For normoxia vs hypoxia experiments, cells were moved to hypoxic incubators (Nuaire NU-5831 for 4% O_2_, Biospherix ProOx C21 for 1% O_2_) 7-8 hours after plating, to ensure that cells had adhered to dish by the time hypoxia treatment started. For the 14-day-long treatments, cells were split every 3 or 4 days and put back in hypoxia once they adhered to the dish. For experiments with small molecule inhibitors, MCF7 or T47D cells were plated, and once cells adhered to the plates, inhibitors were added in fresh media. Inhibitor concentrations and the duration of the treatments are specified in the figure legends. The inhibitors used in this study were FG-4592 (Selleckchem, S1007), GSK-J4 (Selleckchem, S7070), KDM5-C70 (Xcess Biosciences, M60192-2), NSC636819 (Sigma, 5131996), MS275 (Selleckchem, S1053) and UNC0638 (U4885).

All cultures were routinely (every 1-2 months) tested for mycoplasma contamination using MycoAlert (Lonza). Autosomal STR profiles for cell line authentication were performed by the University of Arizona Genetics Core.

### 3D culture

MCF7-HypT cells were FACS sorted for RFP^+^GFP^-^ phenotype. Cells were counted and 20,000 cells were plated in each well of a 24-well ultra-low attachment plate to allow cells to form small aggregates. The following day, Matrigel-collagen mixture was prepared and MCF7-HypT cells were plated into 8-chamber slides (Corning, 354118) following protocol by Nguyen-Ngoc et al (61). Images of spheres were taken the day of plating and every 2-3 days to track the conversion of RFP^+^ cells to GFP^+^ cells over time. Images were taken on a Keyence BZ-9000 fluorescence microscope.

### Generation of stable cell lines

PHAGE2-EF1α-LoxP-DsRED-STOP-LoxP-eGFP (Addgene, 62732) and pLKO.1-5xHRE-CRE-ODD plasmids were used for producing MCF7-HypT cells. To modify the CRE reporter, the bGH poly(A) signal was inserted into the NheI site, downstream of DsRed. The bGH poly(A) signal was PCR amplified from pX459 (Addgene, 62988) using primers forward, 5’-AAGTCTAGACGCTCCCCAGCATGCCTGCT-3’ and reverse, 5’-AGGGCTAGCCGACTGTGCCTTCTAGTTGCCAGCCA-3’. To generate the pLKO-5xHRE-CRE-ODD plasmid, pLKO.1-neo (Addgene, 13425) was used as the backbone. The U6 promoter was deleted and replaced with an XbaI site using NEB Q5 site directed mutagenesis kit (NEB Cat# E0554S) with primers forward, 5’-AAAAATCTAGATGTGGAAAGGACGAAACACC-3’; reverse, 5’-AAAAATCTAGAGAAATAGGCCCTCGGTGAAG-3’ according to manufacturer’s protocol. The reaction was transformed into chemically competent NEB Stable cells (NEB Cat#C3040H) and the plasmid was verified by restriction enzyme digestion and Sanger sequencing. Subsequent cloning steps were performed to generate pLKO-5xHRE-CRE-ODD by using primers forward, 5’-AAAAAACTAGTGCTAGCGGTAGGCGTGTACGGTGGGAGGTCTATATAAGCAGAGCTC – 3’, reverse 5’-AAAAATCTAGAGATCTGACGGTTCACTAAACGAGCTCTGCTTATATAGACC – 3’; forward, 5’-AAAAAGCTAGCTCTAGAACCATGGCCAATTTACTGACCGTACACC-3’, reverse, 5’-AAAAAGCTAGCGGCAACTAGAAGGCACAGTCG – 3’; forward, 5’-AAAAAGCTAGCGGTGGAGAGAGAGACAGAGACAG -3’, reverse, 5’-AAAAAGCTAGCGGTGGATCCTGCAAAAAGAACAAGTAGC -3’, along with the pEF-5xHRE-GFP (Addgene, 46926) and pEF-5xHRE-CRE-ER^T2^-ODD-myc(62) plasmids as templates. For HIF mutant cell lines, we made retrovirus using HA-HIF1α P402A/P564A-pBabe-puro (Addgene, 19005) and HA-HIF2α P405A/P531A/N847Q-pBabe-puro (Addgene, 19006). To generate lentivirus for HIF mutants we subcloned HIF1α mutant from HA-HIF1α P402A/P564A-pcDNA3 (Addgene, 18955) into pCDH-CMV (Addgene, 72265) using BamHI and NotI restriction sites and HIF2α mutant from the pBabe retroviral vector into pCDH-CMV using BamHI and SalI sites.

Lentiviral particles were produced by co-transfecting 293T cells with lentiviral expression vectors in combination with second generation (VSV-G and pCMV-dR.8.91, for pLKO-5xHRE-CRE-ODD, Lenti-dCas9-KRAB-Blast and sgRNA plasmids) or third generation (VSV-G, PMDL and REV, for PHAGE2-EF1α-LoxP-DsRED-STOP-LoxP-eGFP) lentivirus packaging plasmids, using the TransitLT1 transfection reagent (Mirus). Viruses were concentrated with lenti-X concentrator (Clontech). Cell lines to be infected with virus were seeded with 200,000 cells in each well of a 6-well plate. The following day, polybrene (8μg/ml) and concentrated virus were added to each well and incubated overnight. Cells were then selected for infection using puromycin (1 μg/ml), geneticin (0.5 mg/ml) or FACS sorting.

For CRISPR-knockout (KO), guide RNA sequences were cloned into pX458 (Addgene, 48138). sgRNA sequences for ARNT were obtained from previously published paper (Chakraborty et al., 2019): sgRNA1 ARNT, 5’-CACCGGGCTATTAAGCGACGGTCA-3’, 5’-AAACTGACCGTCGCTTAATAGCCC-3’; sgRNA2 ARNT, 5’-CACCGAGAAACGGCCATGCGTAAGA-3’, 5’-AAACTCTTACGCATGGCCGTTTCTC-3’.

Plasmids were transiently transfected into MCF7 cells using Lipofectamine 3000 according to the manufacturer’s protocol (ThermoFisher, L3000015). After three days, GFP+ cells were sorted into individual wells for clonal analysis. Alt-R® Genome Editing Detection Kit (IDT Cat#1075932) was used to screen clones for edited genomes within the targeted region and then validated by western blot for gene knockout.

### Mouse tumor assays

The animal protocol was approved by the Institutional Animal Care and Use Committee of the University of Southern California. For orthotopic injections, 6 to 8 weeks-old female NSG mice (Jackson Laboratory) were anesthetized with isofluorane and GFP-Luc labelled cells or MCF7-HypT cells resuspended in 1:1 PBS and Matrigel (BD Biosciences) were injected into the fourth mammary fat pad. Tumor growth was evaluated by *in vivo* imaging or by measuring the tumor size with a caliper. Tumor volume was calculated using the formula V = π x [d^2^ x D] / 6, were d and D are the minor and major tumor axes respectively. For tail vein injection, 100,000 GFP-Luc expressing cells were injected into the lateral tail vein of 6 to 8 weeks old female NSG mice.

Signals from luciferase-tagged cells were monitored biweekly. Mice were imaged using IVIS Lumina II (Perkin Elmer) following intraperitoneal injection of 150 μl of D-Luciferin substrate at 30 mg/ml (Sid Labs). In all cases, mice were supplemented with 0.36 mg 90-day slow release 17β-Estradiol pellets (Innovative Research of America, Cat# NE-121). For tumor formation experiments with NT2.5 cells, 6 to 8 weeks-old female FVB mice (Jackson Laboratory) were used and tumor cells (1×10^6^) were resuspended in PBS and injected into the fourth mammary fat pad. For full methods describing derivation of NT2.5 tumors used for scRNAseq please see(58).

### Tumor dissociation

Tumors were dissected and collected in cold PBS. A piece of tumor was saved for fixation and immunostaining when needed. Tumors were minced and placed in dissociation media consisting of RPMI supplemented with 2 mg/ml collagenase I, 5% FBS and 15 U/ml DNAse. Samples were dissociated using the gentleMACS dissociator (Miltenyi Biotec), and filtered through a 70 μm cell strainer. After red blood cell lysis, cell suspensions were washed and used as described below.

### Flow cytometry and cell sorting

For immune cells analysis of NT2.5 tumors, dissociated cell suspensions were cryopreserved with 90% FBS and 10% DMSO. Once thawed, 5X10^6^ cells per mL were washed with cold PBS and blocked in PBS with Rat Anti-Mouse CD16/CD32 (Mouse BD Fc Block™, BD Biosciences) for 15 min at 4°C in the dark. Then, staining of cell surface markers for myeloid cell phenotyping was performed by incubating cells with fluorescent-labeled primary antibodies for 30 min at 4 °C in the dark. Cells were then fixed and permeabilized using Cyto-Fast™ Fix/Perm Buffer Set (Biolegend) before intracellular staining of Arginase I. LIVE/DEAD™ Fixable Aqua Dead Cell Stain Kit (ThermoFisher) was used as a viability marker. Antibodies used were CD86 (Biolegend, 105037), F4/80 (Biolegend 123141), MHCII (BD Biosciences 562009), Ly6C (Biolegend, 128012), CD11b (Biolegend, 101207), CD206 (Biolegend, 141732), Ly6G (BD Biosciences, 560599), CD11c (BD Biosciences, 560583), CD3 (BD Biosciences, 560590) and Arginase 1 (Thermo Fisher 48-3697-82). For MHC-I analysis, BRx68 cells grown in different conditions were stained with HLA-ABC antibody (Biolegend, 311410). The myeloid-derived suppressor cells (MDSC) were gated by CD3^-^/CD11^+^/MHCII^-^ and macrophages were gated by CD3^-^/MHCII^+^/F4/80^+^. The granulocytic MDSCs (gMDSCs) were identified as Ly6G^+^/Ly6C^int^ MDSCs and myeloid MDSCs (mMDSCs) were identified as Ly6G^low^/Ly6C^+^ MDSC. The Attune NxT cyometer (Thermo Fisher Scientific) was used for these analyses.

For FACS sorting, cell suspensions from MCF7-HypT tumors were washed with PBS, blocked in PBS with 10% FBS and 1 μg/ml mouse Fc Block (BD Biosciences) for 15 minutes and incubated with the human-specific anti-CD298-APC antibody (Biolegend, 341706) for 30 minutes. DAPI (1 μg/ml) was used as a viability marker. CD298^+^DAPI^-^GFP^+^RFP^-^ and CD298^+^DAPI^-^GFP^-^RFP^+^ cells were sorted into pre-chilled tubes with PBS, centrifuged and lysed with Quick RNA lysis buffer to proceed with RNA extraction. For sorting RFP+ cells from MCF7-HypT cells grown *in vitro*, cells were collected, resuspended in cold 1% BSA in PBS with DAPI (1 μg/ml), and sorted. FACS Aria II (BD Biosciences) was used for cell sorting.

### Single cell library preparation and data processing

The Chromium^TM^ Single Cell Reagent Kit v2 or Next GEM Multiome kit (10x Genomics) were used for single cell RNAseq (Figure S2) or ATAC and gene expression (Figure 6E-F) respectively. On the day of scRNAseq processing, tumors from 12- and 16-week-old mice were dissociated as previously described. After washes with cold PBS with 0.04% BSA, 6000 cells were loaded into a Single Cell A Chip. Tumors processed with Next GEM Multiome kit were fresh frozen prior nuclei isolation. Nuclei isolation and permeabilization was performed following 10x Genomics’ demonstrated protocol (CG000375-RevC). Library preparation was done following manufacturer’s instructions. Libraries were sequenced on NextSeq (Illumina). The raw fastq files were processed with Cell Ranger 3.0.2 (RNAseq) or Cell Ranger ARC 2.0 (Multiome) from 10x Genomics to generate a sparse matrix of features by barcodes. For single cell mutiomic data, alignment was performed using a custom reference package containing annotations for GFP and RFP genes. This output was then loaded into R version 4.0.2 and processed with the R package Seurat 4.0.0(63). The data was then filtered with standard quality control to remove cells with few genes expressed or an overrepresentation of mitochondrial reads. Data was then scaled and normalized. Linear reduction analysis was performed by calculation of PCA from the most variable genes. Cells were then clustered using a resolution of 0.5 and visualized by UMAP. Marker genes were used to identify different cell types present.

IFN/AP and Hypoxia scores were generated using the AddModuleScore function, with a control value of 5. Tumor cells were binned into deciles by IFN/AP score, and Hypoxia score was then plotted for each decile with the DotPlot function. Dropout events in RNA assay from multiomic data were recovered using a method based on low-rank approximation (cite https://www.nature.com/articles/s41467-021-27729-z). Tumor cells were characterized by the expression of RFP or GFP markers and binned into deciles by Hypoxia score. Hypoxic tumor cell were defined as cells in the top Hypoxia decile (top 10%) and moderate hypoxic cells as cells in decile 8 and 9. Then IFN/AP score was plotted for each group with the VlnPlot function and wilcox statistical test was applied using R package ggsignif 0.6.4.

### Fluorescent immunohistochemistry

Primary tumors and lungs were fixed in 4% PFA for 4.5 hours, washed three times with cold PBS and cryoprotected in PBS with 30% sucrose at 4°C overnight. Tissues were then embedded in OCT, frozen using dry ice/ethanol baths and stored at -80°C. Cryosections were incubated at 4°C overnight with antibodies against GFP (Abcam, ab13970), RFP (Abcam, ab62341), pimonidazole (Hypoxyprobe, Pab2627), and CRE (Millipore, mab3120). The secondary antibodies used in this study were Goat anti-Chicken IgY Alexa Fluor 488 (Life Technologies, A11039), Goat anti-Rabbit IgG Alexa Fluor 594 (Life Technologies, A11012) and Goat anti-mouse IgG Alexa Fluor 647 (Life Technologies, A21240). All secondary antibodies were incubated for one hour at room temperature. Images were taken on a Keyence BZ-9000 or an Axioimager A1 (Zeiss) fluorescence microscope.

### Gene expression profiling

Integrity of RNA extracted with Quick-RNA MicroPrep (Zymo) was measured using Bioanalyzer (Agilent). Sequencing libraries were prepared with KAPA stranded RNA-Seq Kit with RiboErase (Kapa Biosystems) according to manufacturer’s instructions, and single-end 75 bp sequencing was performed at the USC Molecular Genomics Core, using a NextSeq high output 75 cycle kit (Illumina). Reads were trimmed for Nextera and Illumina adapter sequences using trimgalore(64) with default parameters. Trimmed reads were mapped to hg19 (GRCh37) reference using STAR (v2.7.1a)(65), and the reads mapped to the genes annotated in the ENSEMBL GRCh37.p13 GTF (release 75) were quantified using HTSeq-count (v0.11.2)(66). Differentially expressed genes across hypoxia-normoxia conditions were identified after controlling for cell line and batch effects (FDR≤0.05), using DESeq2 (v1.26.0)(67). PCA and heatmaps were generated using variance stabilizing transformed normalized counts after regressing out the batch effects using removeBatchEffect function from limma package (v3.42.2)(68). Pathway analysis was performed using Ingenuity Pathway Analysis (QIAGEN, https://www.qiagenbioinformatics.com/products/ingenuitypathway-analysis) and overlapping gene sets were found using Gene Set Enrichment Analysis at https://www.gsea-msigdb.org/gsea(69,70).

RNA-Seq data of different breast cancer cell lines were obtained from Gene Expression Omnibus (GEO). Genes differentially expressed in responder cell lines (MCF7, T47D, and BT4774) compared to non-responder cell lines (231 and HS578T) were identified using DESeq2 after controlling for cell lines differences and study specific difference at their respective GSE level. Microarray expression data of mouse Py117 and Py8119 cancer cells were obtained from (GSE118181)(71). Differentially expressed (DE) genes between responsive and non-responsive clones were identified using limma(68) at an FDR level of 0.05. After mapping RefSeq IDs to ENSEMBL IDs, common orthologous DE genes in responder RNA-seq breast cancer cell lines and mouse Py117 were identified from the two analysis.

### qRT-PCR

Total RNA extracted with Quick-RNA MicroPrep (Zymo) was reverse transcribed using iScript Reverse Transcription Supermix (Bio-Rad), according to the manufacturer’s instructions. Real time PCR was performed on a CFX96^TM^ Real-Time PCR detection system (Bio-Rad), using iTaq^TM^ Universal SYBR® Green Supermix (Bio-Rad). The primer sequences used were human IFIT1 (forward, 5’-AGAAGCAGGCAATCACAGAAAA-3’; reverse, 5‘-CTGAAACCGACCATAGTGGAAAT-3’); human IFIT3 (forward, 5’-‘AGGCTCTGGAAGAAAAGCCC-3’, reverse, 5’-TTCTGCAGTTTCAGGCCCAA-3’); human ISG15 (forward, 5-‘GTACAGGAGCTTGTGCCGTG-3’; reverse, 5’-ACACCTGGAATTCGTTGCCC-3’); human OAS1 (forward, 5’-TGTCCAAGGTGGTAAAGGGTG-3’; reverse, 5’-CCGGCGATTTAACTGATCCTG-3’ human CA9 (forward, 5’-GGGTGTCATCTGGACTGTGTT-3’; reverse, 5’-CTTCTGTGCTGCCTTCTCATC-3’); human PHD3 (forward, 5’-GGCTGGGCAAATACTACGTCAA-3’; reverse, 5’-CCTGTTCCATTTCCCGGATAG-3’); human NDRG1 (forward, 5’-GTCCTTATCAACGTGAACCCTT-3’; reverse, 5’-GCATTGGTCGCTCAATCTCCA-3’); human ESR1 (forward, 5’-GTGGGATACGAAAAGACCGA-3’; reverse, 5’-AAGGTTGGCAGCTCTCATGT-3’); human PS2 (forward, 5’-TCGGGGTCGCCTTTGGAGCAG-3’; reverse, 5’-GAGGGCGTGACACCAGGAAAACCA-3’); human 36B4 (forward, 5’-GTGTTCGACAATGGCAGCAT-3’; reverse, 5’-AGACACTGGCAACATTGCGGA-3’); mouse ifit1 (forward, 5’-TCTGCTCTGCTGAAAACCCAG-3’; reverse, 5’-CAAGGAACTGGACCTGCTCTG-3’); mouse isg15 (forward, 5’-GAGCTAGAGCCTGCAGCAAT-3’; reverse, 5’-TCACGGACACCAGGAAATCG-3’); mouse ca9 (forward, 5’-TCTGAAGACAGGATGGAGGAGT-3’; reverse, 5’-AGAGTAGGGTGCCTCCATAG-3’); mouse phd3 (forward, 5’-GGGACAGGTTATGTTCGCCA-3’; reverse, 5’-TGCGGTCTGACCAGAAGAAC-3’); mouse 36B4 (forward, 5’-CTCGTTGGAGTGACATCGTCT-3’; reverse, 5’-GTCTGCTCCCACAATGAAGC-3’). The relative RNA levels were calculated using the DDCq method, using 36B4 as reference gene.

### Western blotting

Cells were washed with PBS and lysed in Laemmli buffer (50 mM Tris, pH6.8, 1.25% sodium dodecyl sulfate, 15% glycerol). Cell lysates were heated at 95°C for 15 minutes for complete lysis and denaturalization, after which proteins were quantified by Lowry protein assay (Bio-Rad) and 5% (v/v) β-mercaptoethanol (Sigma) and 0.004% (w/v) bromophenol-blue (Sigma) were added. Proteins were run on 4-15% SDS-polyacrylamide gradient gels (Bio-Rad) and transferred to a low-fluorescence PVDF membrane (Millipore). Primary antibodies used in this study are HIF1α (BD Biosciences, 610958), HIF2α (R&D, AF2886), ARNT (Cell Signaling, 5537), H3 (Abcam, ab1791), H3K4me3, (Epigentek, A-4033), H3K9me2 (Abcam, ab1220), H3K9me3 (Epigentek, A-4036), H3K27me3 (Millipore, 07-449), H3K36me3 (Epigentek, A4042), H3K27ac (Active Motif, 39133), IRF3 (Cell Signaling, 10949), FOXO1 (Cell Signaling, 2880), FOXA1 (Abcam, ab23738), FOXO3a (Cell Signaling, 2497), and β-actin (Sigma, A5441). Membranes were probed at 4°C overnight, followed by incubation with an IRDye®-conjugated anti-mouse (LI-COR, 926-32210), anti-rabbit (LI-COR, 926-68071) or anti-goat secondary (LI-COR, 926-32214) antibodies and images were acquired with a LI-COR Odyssey CLx Near-Infrared Fluorescence Imaging System. Alternatively, anti-mouse-HRP (BioRad, 1706516) or anti-rabbit-HRP (Jackson Labs, 711035152) antibodies were used.

### Statistical analysis

Results represent mean±SEM, unless otherwise indicated. Statistical significance between two groups was determined by unpaired Student’s t-test or Mann-Whitney test, or ANOVA for more than 2 groups followed by Fisher’s LSD test. Statistical analysis was done with GraphPad Prism v.9.

