## Supplemental figures and tables for "Hypoxic memory of tumor intrinsic type I interferon suppression promotes breast cancer metastasis"

### FIGURE S1

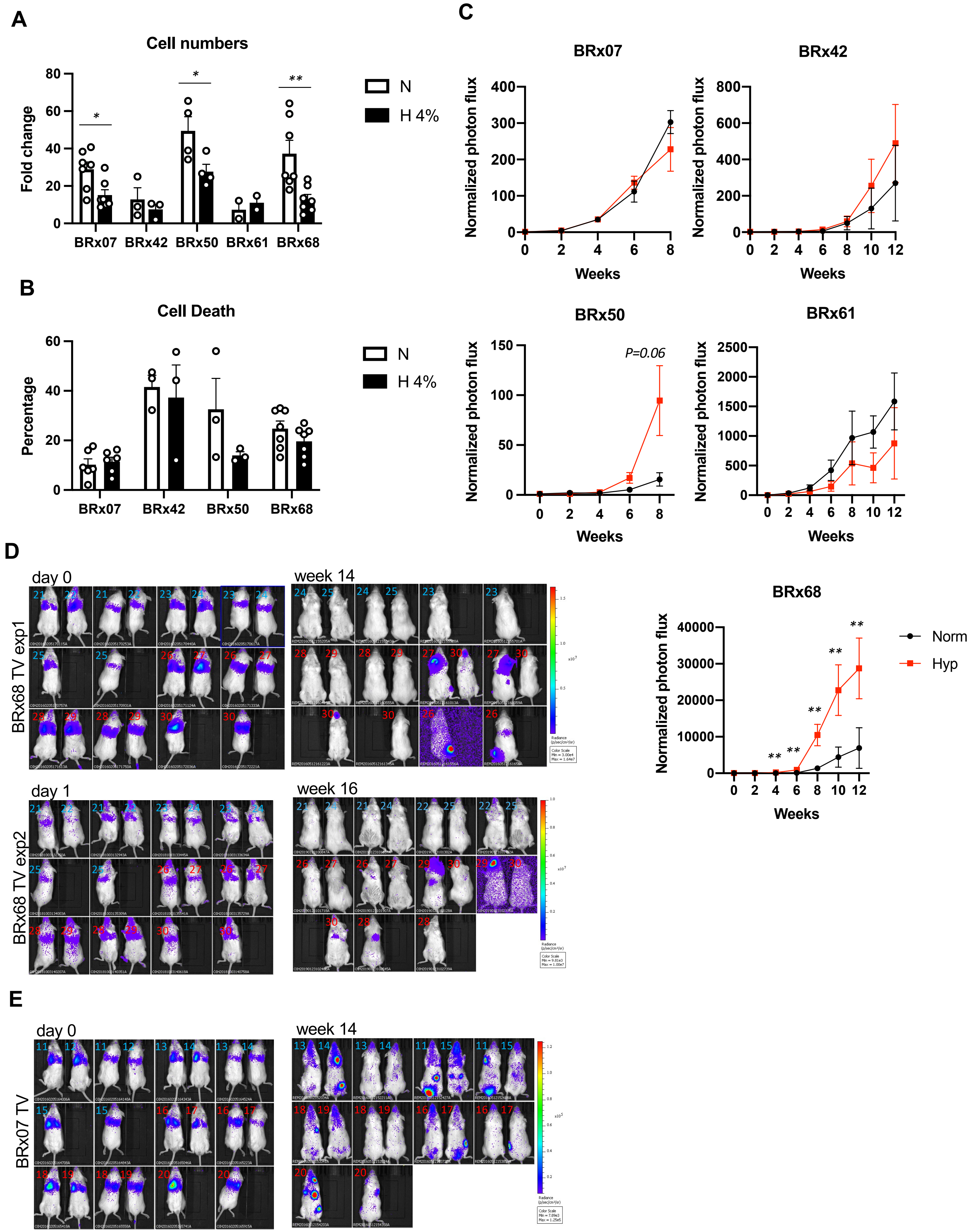
A
Cell numbers
Fold change
80
60
40
20
0
*
*
**
BRx07 BRx42 BRx50 BRx61 BRx68
N
H 4%
B
Cell Death
Percentage
60
40
20
0
BRx07 BRx42 BRx50 BRx68
N
H 4%
C
BRx07
Normalized photon flux
400
300
200
100
0
0 2 4 6 8
Weeks
BRx42
Normalized photon flux
800
600
400
200
0
0 2 4 6 8 10 12
Weeks
BRx50
P=0.06
Normalized photon flux
150
100
50
0
0 2 4 6 8
Weeks
BRx61
Normalized photon flux
2500
2000
1500
1000
500
0
0 2 4 6 8 10 12
Weeks
D
day 0
week 14
BRx68 TV exp1
day 1
week 16
BRx68 TV exp2
BRx68
Normalized photon flux
40000
30000
20000
10000
0
0 2 4 6 8 10 12
Weeks
Norm
Hyp
**
E
day 0
week 14
BRx07 TV

**Figure S1:** ***In vitro*** **and** ***in vivo*** **growth of CTC lines influenced by different oxygen culture conditions.**

**(A)** Fold change in cell numbers after 8 weeks growing in normoxia or hypoxia. Mean±SEM. *P<0.05, **P<0.01. P value was calculated with two-tailed Student's *t* test. **(B)** Percentage death cells in CTCs grown in normoxia or hypoxia for 8 weeks. Mean±SEM. * p<0.05, ** p<0.01. P value was calculated with two-tailed Student's *t* test. **(C)** Biweekly bioluminescence values from mice orthotopically injected with 2,000 cells per mouse for BRx07, BRx61 and BRx68, 20,000 cells for BRx42 and BRx50. Prior to the injection, cells were grown in normoxia or hypoxia for 8 weeks. For each timepoint, the bioluminescence is normalized with the photon flux of each mouse on the day of injection. Mean±SEM. *P<0.05, **P<0.01. P value was calculated with Welch's t test. **(D-E)** Bioluminescence imaging data of mice injected with 100,000 BRx68-GFP/luc **(D)** or BRx07-GFP/luc **(E)** cells. Images taken on the day of the injection and 14 weeks after the injection are shown (both front and back).

### FIGURE S2

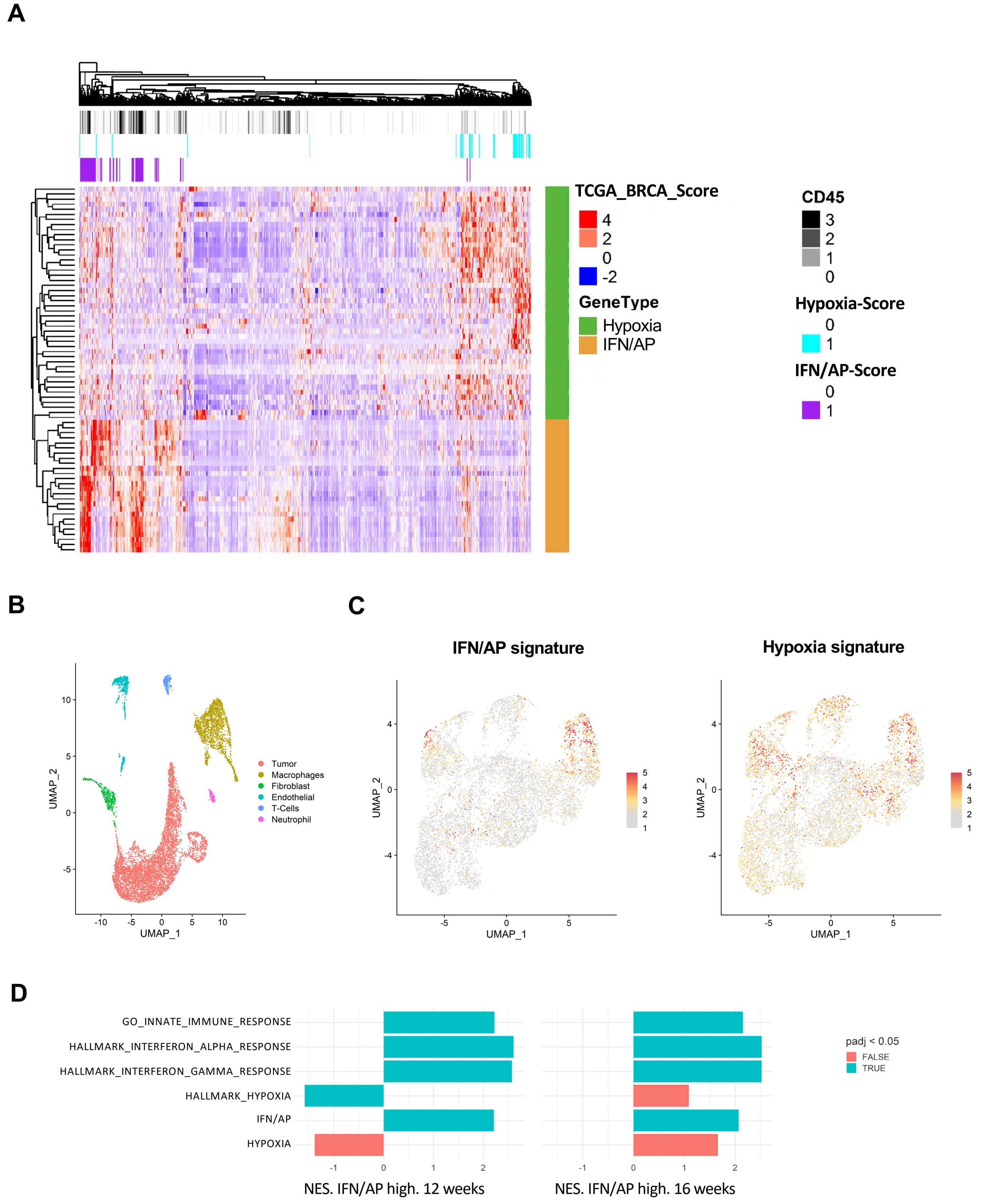
A
TCGA_BRCA_Score
4
2
0
-2
GeneType
Hypoxia
IFN/AP
CD45
3
2
1
0
Hypoxia-Score
0
1
IFN/AP-Score
0
1
B
UMAP_2
UMAP_1
Tumor
Macrophages
Fibroblast
Endothelial
T-Cells
Neutrophil
C
IFN/AP signature
Hypoxia signature
UMAP_2
UMAP_1
D
GO_INNATE_IMMUNE_RESPONSE
HALLMARK_INTERFERON_ALPHA_RESPONSE
HALLMARK_INTERFERON_GAMMA_RESPONSE
HALLMARK_HYPOXIA
IFN/AP
HYPOXIA
NES. IFN/AP high. 12 weeks
NES. IFN/AP high. 16 weeks
padj < 0.05
FALSE
TRUE

**Figure S2: *In vivo* breast tumors show inverse correlation between hypoxia and IFN/AP gene signatures.**

**(A)** Heatmap of hypoxia and IFN/AP signatures from RNAseq data of breast tumors in TCGA. **(B)** UMAP plot showing the cell type clusters from all 4 tumors (2 at 12 weeks and 16 weeks each). **(C)** UMAP plot showing IFN/AP and hypoxia signatures in tumor cells only. **(D)** GSEA analysis of enriched interferon and hypoxia pathways in IFN/AP high cells at 12 weeks (left) and 16 weeks (right).

### FIGURE S3

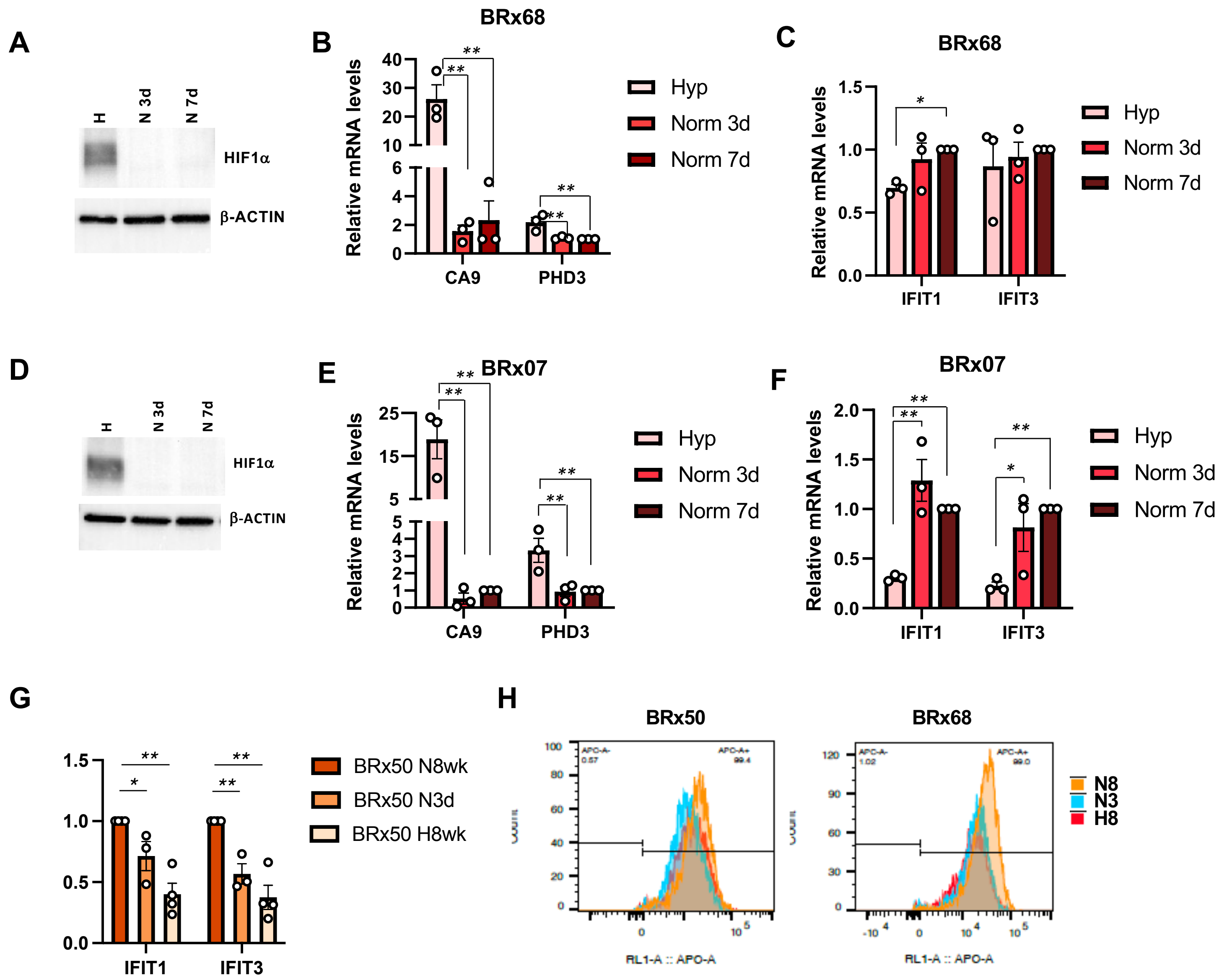
A
H
N 3d
N 7d
HIF1α
β-ACTIN
B
BRx68
Relative mRNA levels
CA9
PHD3
Hyp
Norm 3d
Norm 7d
C
BRx68
Relative mRNA levels
IFIT1
IFIT3
Hyp
Norm 3d
Norm 7d
D
H
N 3d
N 7d
HIF1α
β-ACTIN
E
BRx07
Relative mRNA levels
CA9
PHD3
Hyp
Norm 3d
Norm 7d
F
BRx07
Relative mRNA levels
IFIT1
IFIT3
Hyp
Norm 3d
Norm 7d
G
IFIT1
IFIT3
BRx50 N8wk
BRx50 N3d
BRx50 H8wk
H
BRx50
BRx68
RL1-A :: APO-A
N8
N3
H8

**Figure S3: Hypoxia suppression of IFN/AP lasts longer than the hypoxic exposure.**

**(A-C)** Immunoblot analysis with antibodies against HIF1α and β-ACTIN (**A**) and qPCR analysis of mRNA expression levels of hypoxia targets (**B**) and IFN targets (**C**) in BRx68 cells reoxygenated for 3 or 7 days. Mean±SEM, n=3. P values were calculated with one-way ANOVA followed by Fisher' LSD test. $^{*}P<0.05$, $^{**}P<0.01$. **(D-F)** Immunoblot analysis with antibodies against HIF1α and β-ACTIN (**D**) and qPCR analysis of mRNA expression levels of hypoxia targets (**E**) and IFN targets (**F**) in BRx07 cells reoxygenated for 3 or 7 days. Mean±SEM, n=3. P values were calculated with one-way ANOVA followed by Fisher' LSD test. $^{*}P<0.05$, $^{**}P<0.01$. **(G)** qPCR analysis of IFN target gene mRNA expression levels in BRx50 cells grown in hypoxia for 8 weeks (4% O2, H8) or normoxia for 8 weeks (N8) or 3 days (N3). Mean ±SEM, n=5 (IFIT1) and n=3 (IFIT3). P values were obtained with one-way ANOVA followed by Fisher's LSD test. $^{*}P<0.05$, $^{**}P<0.01$. **(H)** Histogram for FACS analysis of HLA-ABC levels for BRx50 and BRx68 cells grown in hypoxia for 8 weeks (4% O2, H8) or normoxia for 8 weeks (N8) or 3 days (N3).

FIGURE S4

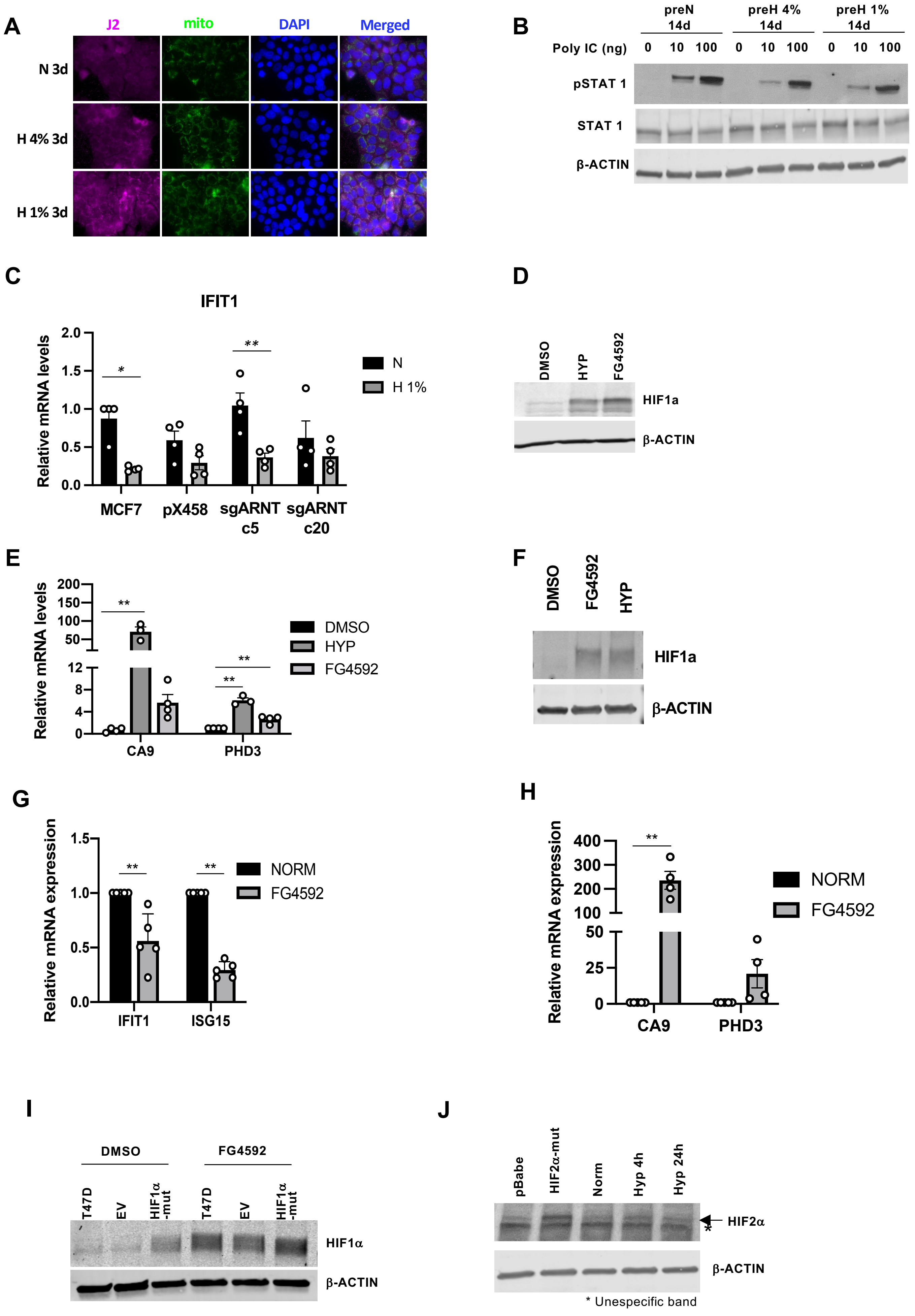
A
J2
mito
DAPI
Merged
N 3d
H 4% 3d
H 1% 3d
B
preN 14d
preH 4% 14d
preH 1% 14d
Poly IC (ng)
0 10 100 0 10 100 0 10 100
pSTAT 1
STAT 1
β-ACTIN
C
IFIT1
Relative mRNA levels
2.0
1.5
1.0
0.5
0.0
*
**
N
H 1%
MCF7
pX458
sgARNT c5
sgARNT c20
D
DMSO
HYP
FG4592
HIF1a
β-ACTIN
E
Relative mRNA levels
200
150
100
50
12
8
4
0
**
**
**
DMSO
HYP
FG4592
CA9
PHD3
F
DMSO
FG4592
HYP
HIF1a
β-ACTIN
G
Relative mRNA expression
1.5
1.0
0.5
0.0
**
**
NORM
FG4592
IFIT1
ISG15
H
Relative mRNA expression
400
300
200
100
50
25
0
**
NORM
FG4592
CA9
PHD3
I
DMSO
FG4592
T47D
EV
HIF1α -mut
T47D
EV
HIF1α -mut
HIF1α
β-ACTIN
J
pBabe
HIF2α-mut
Norm
Hyp 4h
Hyp 24h
HIF2α
*
β-ACTIN
* Unespecific band

**Figure S4: Evaluation of stimulus, response, and HIF for hypoxic downregulation of IFN/AP signal.**

**(A)** Immunofluorescence staining with antibodies against J2 (dsRNA) and mitochondria in MCF7 cells cultured in normoxia, 4% or 1% hypoxia for 3 days. Nuclei are stained with DAPI. **(B)** Immunoblot analysis of pSTAT1, STAT1, and β-actin levels in MCF7 cells transfected with 0, 10, and 100ng of poly IC, previously cultured in normoxia, or 4% or 1% $O_2$ condition for 14 days. **(C)** Relative mRNA expression levels of IFIT1 in bulk MCF7 cells, 1 px458 control MCF7 clone and 2 ARNT-KO MCF7 clones grown in normoxia or hypoxia (1% $O_2$) for 3 days. Mean±SEM, n=4. P values were obtained with one-way ANOVA followed by Fisher's LSD test. *P<0.05, **P<0.01. **(D-E)** Immunoblot analysis with antibodies against HIF1α and β-ACTIN **(D)** and qPCR analysis of CA9 and PHD3 expression levels **(E)** in MCF7 cells treated with 100 μM FG4592 or hypoxia for 3 days. Mean±SEM, n=4. P values were obtained with one-way ANOVA followed by Fisher's LSD test. *P<0.05, **P<0.01. **(F-H)** Immunoblot analysis with antibodies against HIF1α and β-ACTIN **(F)** and qPCR analysis of CA9 and PHD3 **(G)** and IFIT1 and ISG15 **(H)** expression levels in T47D cells treated with 100 μM FG4592 or hypoxia for 3 days. Mean±SEM, n=5. P values were obtained 2-tailed Student's t test. **P<0.01 **(I)** Immunoblot analysis of HIF1α and β-ACTIN levels in T47D cells, and T47D with expression of EV, or HIF1α-mutant with DMSO or FG4592 treatment in normoxia. (J) Immunoblot analysis of HIF2α and β-ACTIN levels in T47D cells expressing pBabe-EV, or HIF2α-mutant cells cultured in normoxia or hypoxia. *denotes a non-specific band.

### FIGURE S5

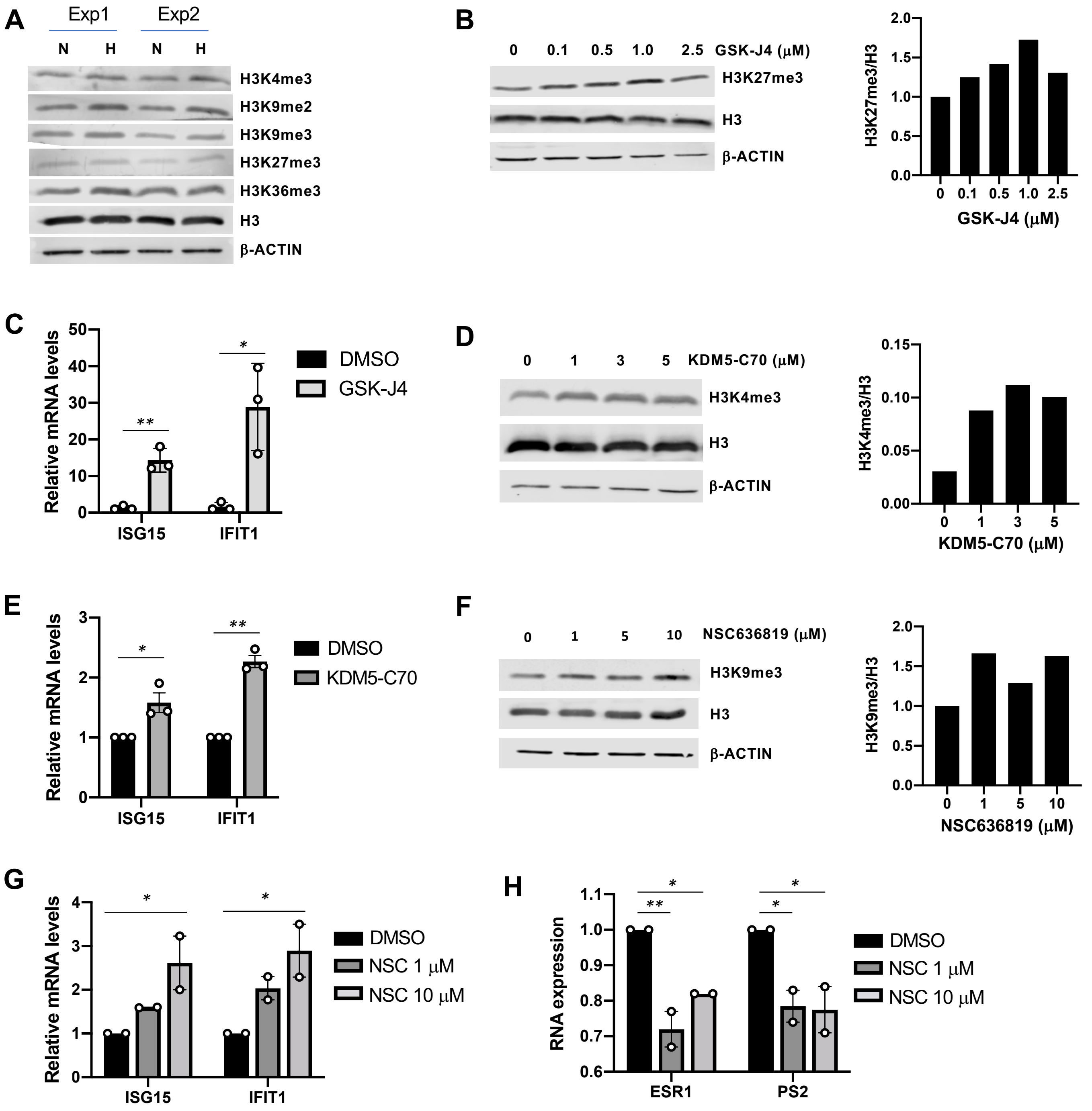

A
Exp1
Exp2
N H N H
H3K4me3
H3K9me2
H3K9me3
H3K27me3
H3K36me3
H3
β-ACTIN
B
0 0.1 0.5 1.0 2.5 GSK-J4 (μM)
H3K27me3
H3
β-ACTIN
H3K27me3/H3
GSK-J4 (μM)
C
Relative mRNA levels
DMSO
GSK-J4
ISG15
IFIT1
D
0 1 3 5 KDM5-C70 (μM)
H3K4me3
H3
β-ACTIN
H3K4me3/H3
KDM5-C70 (μM)
E
Relative mRNA levels
DMSO
KDM5-C70
ISG15
IFIT1
F
0 1 5 10 NSC636819 (μM)
H3K9me3
H3
β-ACTIN
H3K9me3/H3
NSC636819 (μM)
G
Relative mRNA levels
DMSO
NSC 1 μM
NSC 10 μM
ISG15
IFIT1
H
RNA expression
DMSO
NSC 1 μM
NSC 10 μM
ESR1
PS2

**Figure S5. Evaluation of epigenetic reasons for hypoxic downregulation of IFN/AP signal. (A)** Immunoblot analysis showing changes in different H3 histone modifications in MCF7 cells grown in normoxia or hypoxia (1% $O_2$). **(B)** Immunoblot analysis with antibodies against H3K27me3, H3 and β-actin in MCF7 cells treated with different concentrations of GSK-J4. Graphs on the right show quantification of the signal. **(C)** qPCR analysis of ISG15 and IFIT1 expression levels in MCF7 cells treated with 1 mM GSK-J4 (KDM6 inhibitor). P values were obtained with 2-tailed Student's t test. *P<0.05, **P<0.01. **(D)** Immunoblot analysis with antibodies against H3K4me3, H3 and β-ACTIN in MCF7 cells treated with different concentrations of KDM5-C70. Graph on the right shows corresponding signal quantification. **(E)** qPCR analysis of ISG15 and IFIT1 expression levels in MCF7 cells treated with 3 mM KDM5-C70 (KDM5 inhibitor). P values were obtained with 2-tailed Student's t test. *P<0.05, **P<0.01. **(F)** Immunoblot analysis with antibodies against H3K9me3, H3 and β-ACTIN in MCF7 cells treated with different concentrations of NSC636819. Graph on the right shows corresponding signal quantification. **(G)** qPCR analysis of ISG15 and IFIT1 expression levels in MCF7 cells treated with 1 or 5 mM NSC636819 (KDM4 inhibitor). **(H)** qPCR analysis of ESR1 and PS2 expression levels in MCF7 cells treated with 1 or 5 mM NSC636819. mean±SEM. Statistical significance was calculated with one-way ANOVA followed by Fisher's LSD test. *P<0.05, ** P<0.01.

#### FIGURE S6

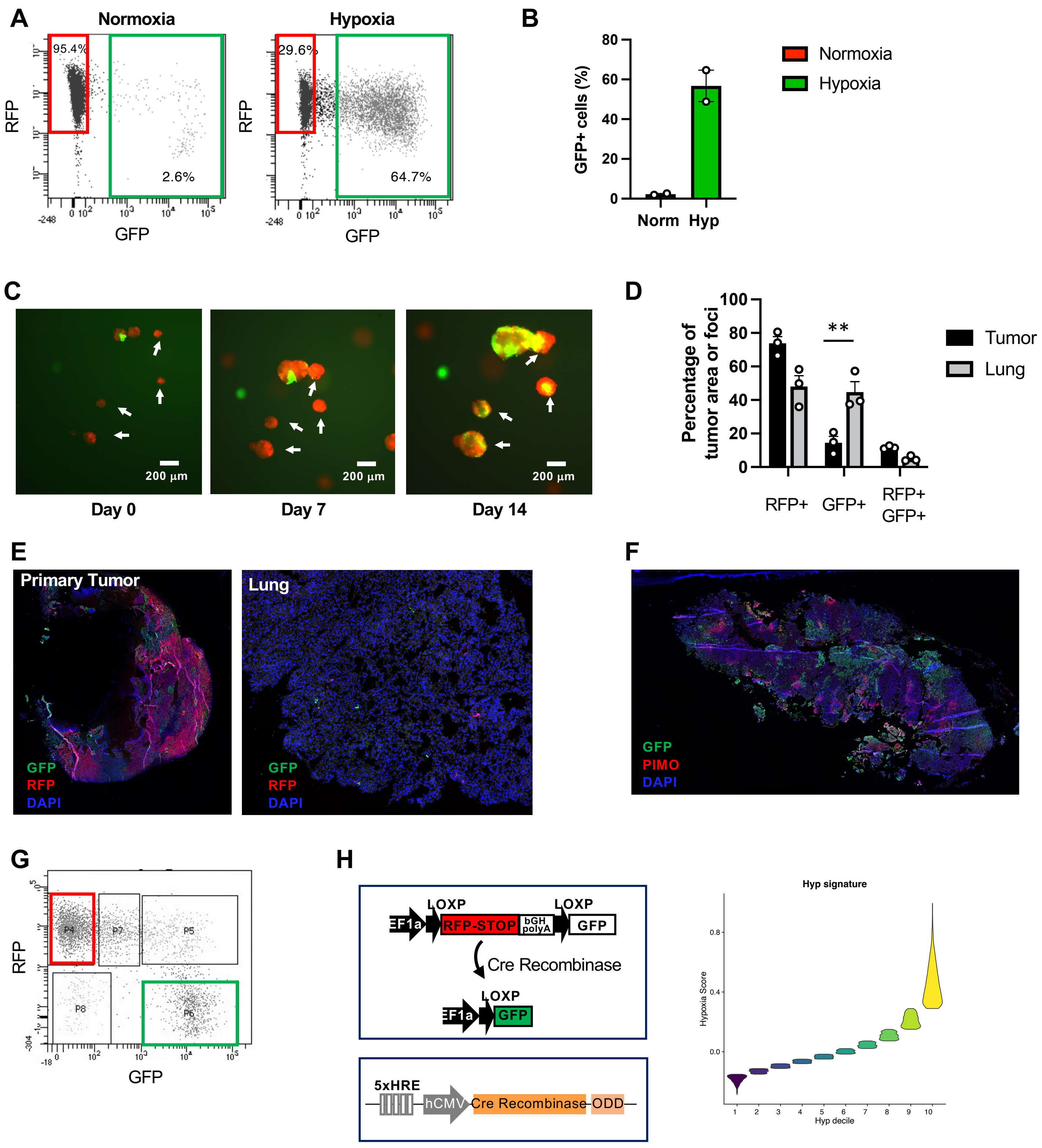

A
Normoxia
Hypoxia
95.4%
2.6%
29.6%
64.7%
RFP
GFP
B
GFP+ cells (%)
Norm
Hyp
Normoxia
Hypoxia
C
200 μm
Day 0
Day 7
Day 14
D
Percentage of tumor area or foci
Tumor
Lung
**
RFP+
GFP+
RFP+ GFP+
E
Primary Tumor
Lung
GFP
RFP
DAPI
F
GFP
PIMO
DAPI
G
P4
P7
P5
P8
P6
RFP
GFP
H
LOXP
EF1a
RFP-STOP
bGH polyA
GFP
Cre Recombinase
5xHRE
hCMV
Cre Recombinase
ODD
Hyp signature
Hypoxia Score
Hyp decile

**Figure S6: Hypoxia-tracing reporter system showed tumor cells with hypoxic memory and enhanced metastatic capacity.**

**(A-B)** Representative FACS plots **(A)** and bar graph **(B)** showing the percentage of GFP+ cells in MCF7-HypT cells cultured in normoxia or hypoxia (1% $O_2$) for 3 days. **(C)** Fluorescence microscopy images of MCF7-HypT cells cultured in 3D collagen-Matrigel gels. Arrows point at spheres formed exclusively by RFP-positive cells on the day of starting the 3D culture. Scale bar = 200 mm. **(D)** Bar graph showing quantification of percentage of RFP, GFP or double positive area in primary tumors and RFP, GFP or double positive metastatic foci in the lung of the MCF7-HypT mice. Means ±SEM, n=3. P values were obtained with one-way ANOVA followed by Fisher's LSD test. ** P<0.01. **(E)** Representative example of a primary tumor section, and a lung section from a mouse injected with MCF7-HypT cells. **(F)** Representative example of a tumor stained with antibodies against pimonidazole (red) and GFP (green). DAPI (blue) was used to stained nuclei. **(G)** Representative FACS plot showing GFP and RFP expression in tumor cells isolated from MCF7-HypT tumors. **(H)** Schematic showing the optimized HypT plasmid with addition of 3'UTR of RFP. **(I)** Decile plot showing combined RFP- or GFP-transcript positive cells ranked with hypoxia score.

### FIGURE S7

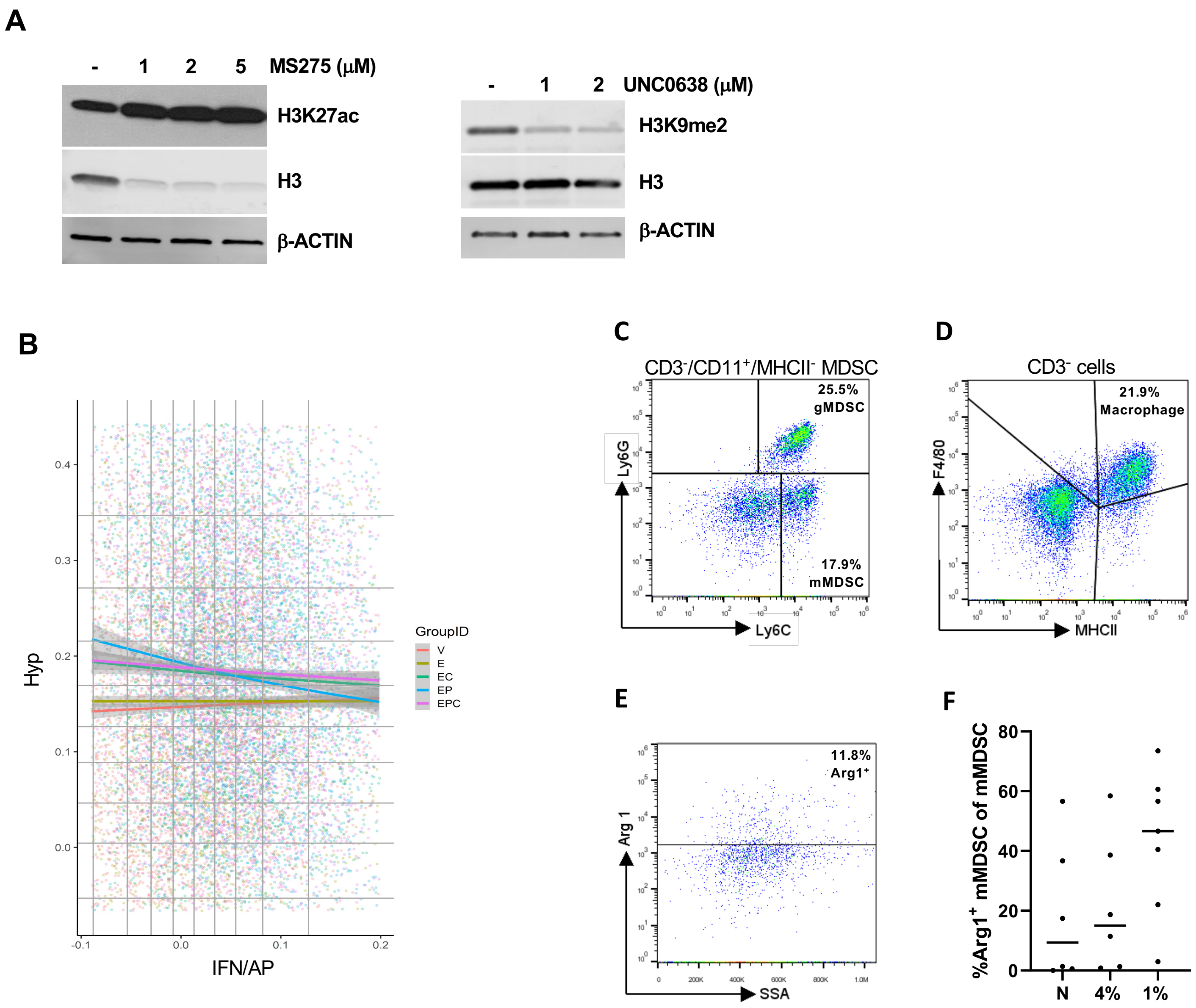
A
- 1 2 5 MS275 (μM)
H3K27ac
H3
β-ACTIN
- 1 2 UNC0638 (μM)
H3K9me2
H3
β-ACTIN
B
GroupID
V
E
EC
EP
EPC
Hyp
IFN/AP
C
CD3-/CD11+/MHCII- MDSC
25.5%
gMDSC
17.9%
mMDSC
Ly6G
Ly6C
D
CD3- cells
21.9%
Macrophage
F4/80
MHCII
E
11.8%
Arg1+
Arg 1
SSA
F
%Arg1+ mMDSC of mMDSC
N
4%
1%

**Figure S7: Hypoxic memory of tumor cells and immune profiles of NT2.5 syngeneic tumors.** **(A)** Immunoblot analysis of H3K27ac (left), H3K9me2 (right) and total H3 levels in MCF7 cells treated with different concentrations of MS275 or UNC0638. β-ACTIN is shown as a loading control. **(B)** Dot plot showing IFN/AP (x-axis) and hypoxia (y-axis) scores for tumor cells (within 5-95 percentiles) for all treatment conditions (as in Fig. 4d). **(C-E)** Representative FACS plots for analysis of gMDSC and mMDSC **(C)**, macrophages **(D)** and $Arg1^+$ cells **(E)**. **(F)** Graph showing percentage of Arg+ mMDSCs in the tumors formed by NT2.5 cells cultured in normoxia, or 4% or 1% hypoxia conditions for 14 days.

**Table S1:** List of genes downregulated in BRx68 cells grown in hypoxia for 8 weeks (≥1.5 Fold change, FDR≤0.05).

**Table S2:** List of genes upregulated in BRx68 cells grown in hypoxia for 8 weeks (≥1.5 Fold change, FDR≤0.05).

**Table S3:** GSEA analysis with genes downregulated in 68H compared to 68N.

**Table S4:** GSEA analysis with genes upregulated in 68H compared to 68N.

**Table S5:** GSEA analysis with genes upregulated in responders compared to non-responders.

**Table S6:** GSEA analysis with genes downregulated in responders compared to non-responders.

**Table S7:** Genes belonging to the hypoxia and IFN/AP signatures

### TABLE S1

Down in 68H (241)

| ensembl_gene_id | gene_symbol | log2FoldChange | pvalue | padj |
|---|---|---|---|---|
| ENSG00000196141 | SPATS2L | -1.776498 | 3.33E-14 | 4.93E-10 |
| ENSG00000179583 | CIITA | -2.6283428 | 7.77E-14 | 5.76E-10 |
| ENSG00000119917 | IFIT3 | -2.7602017 | 1.18E-13 | 5.86E-10 |
| ENSG00000111052 | LIN7A | -1.9774747 | 1.79E-13 | 6.63E-10 |
| ENSG00000101347 | SAMHD1 | -1.9015404 | 3.41E-13 | 1.01E-09 |
| ENSG00000188064 | WNT7B | -1.4015254 | 1.10E-12 | 2.73E-09 |
| ENSG00000183486 | MX2 | -2.5347069 | 4.72E-12 | 1.00E-08 |
| ENSG00000120251 | GRIA2 | -3.8236602 | 9.69E-12 | 1.60E-08 |
| ENSG00000185880 | TRIM69 | -1.5264816 | 1.72E-11 | 2.55E-08 |
| ENSG00000111335 | OAS2 | -1.6163957 | 1.71E-10 | 2.31E-07 |
| ENSG00000168016 | TRANK1 | -1.6560549 | 2.48E-10 | 3.07E-07 |
| ENSG00000133477 | FAM83F | -1.3509354 | 5.99E-10 | 6.83E-07 |
| ENSG00000137752 | CASP1 | -2.1035111 | 9.16E-10 | 9.11E-07 |
| ENSG00000184371 | CSF1 | -2.3315247 | 1.50E-09 | 1.39E-06 |
| ENSG00000185745 | IFIT1 | -2.0654794 | 1.96E-09 | 1.61E-06 |
| ENSG00000115415 | STAT1 | -1.3394175 | 1.92E-09 | 1.61E-06 |
| ENSG00000102699 | PARP4 | -2.3538427 | 2.75E-09 | 2.15E-06 |
| ENSG00000132274 | TRIM22 | -3.0373212 | 3.40E-09 | 2.39E-06 |
| ENSG00000205002 | AARD | -1.6662991 | 3.55E-09 | 2.39E-06 |
| ENSG00000070190 | DAPP1 | -1.2898204 | 3.49E-09 | 2.39E-06 |
| ENSG00000196684 | HSH2D | -1.6195473 | 6.42E-09 | 4.14E-06 |
| ENSG00000132530 | XAF1 | -1.9124961 | 1.13E-08 | 6.88E-06 |
| ENSG00000105855 | ITGB8 | -1.5881034 | 1.16E-08 | 6.88E-06 |
| ENSG00000180263 | FGD6 | -1.2978043 | 1.48E-08 | 8.42E-06 |
| ENSG00000101298 | SNPH | -1.6678123 | 2.36E-08 | 1.29E-05 |
| ENSG00000173334 | TRIB1 | -1.1962519 | 2.54E-08 | 1.34E-05 |
| ENSG00000187608 | ISG15 | -2.2993246 | 2.86E-08 | 1.44E-05 |
| ENSG00000101384 | JAG1 | -1.6934557 | 2.91E-08 | 1.44E-05 |
| ENSG00000168062 | BATF2 | -3.2834633 | 3.51E-08 | 1.68E-05 |
| ENSG00000116096 | SPR | -1.3313656 | 4.22E-08 | 1.96E-05 |
| ENSG00000064763 | FAR2 | -1.6300769 | 4.41E-08 | 1.98E-05 |
| ENSG00000115267 | IFIH1 | -1.4679117 | 5.09E-08 | 2.22E-05 |
| ENSG00000123240 | OPTN | -1.2178157 | 7.41E-08 | 3.06E-05 |
| ENSG00000151726 | ACSL1 | -1.1379393 | 8.26E-08 | 3.23E-05 |
| ENSG00000177409 | SAMD9L | -1.5531509 | 8.80E-08 | 3.35E-05 |
| ENSG00000140853 | NLRC5 | -1.8564224 | 1.08E-07 | 4.01E-05 |
| ENSG00000197461 | PDGFA | -1.7088863 | 1.18E-07 | 4.27E-05 |
| ENSG00000003400 | CASP10 | -2.1799015 | 1.45E-07 | 4.97E-05 |
| ENSG00000182621 | PLCB1 | -1.5994322 | 1.51E-07 | 4.97E-05 |
| ENSG00000164054 | SHISA5 | -1.0560022 | 3.26E-07 | 0.00010066 |
| ENSG00000205413 | SAMD9 | -1.6832595 | 3.36E-07 | 0.00010168 |
| ENSG00000174498 | IGDCC3 | -2.9966538 | 3.89E-07 | 0.00011301 |
| ENSG00000089127 | OAS1 | -1.4639431 | 4.01E-07 | 0.00011441 |
| ENSG00000105357 | MYH14 | -0.9907092 | 4.11E-07 | 0.00011492 |
| ENSG00000157601 | MX1 | -1.5150469 | 4.28E-07 | 0.00011765 |
| ENSG00000152778 | IFIT5 | -1.1669522 | 5.26E-07 | 0.0001418 |
| ENSG00000155846 | PPARGC1B | -1.6440017 | 7.04E-07 | 0.00017998 |
| ENSG00000105953 | OGDH | -1.3426455 | 1.50E-06 | 0.00035855 |
| ENSG00000002549 | LAP3 | -1.1014246 | 1.56E-06 | 0.00036118 |
| ENSG00000221963 | APOL6 | -1.1909711 | 1.97E-06 | 0.00042998 |
| ENSG00000143390 | RFX5 | -0.8187269 | 1.97E-06 | 0.00042998 |
| ENSG00000148346 | LCN2 | -1.1812073 | 2.21E-06 | 0.00047522 |
| ENSG00000112343 | TRIM38 | -0.916375 | 2.45E-06 | 0.00051981 |
| ENSG00000130487 | KLHDC7B | -4.061847 | 2.57E-06 | 0.00052991 |

### TABLE S1 (continued)

| | | | | |
|---|---|---|---|---|
| ENSG00000140464 | PML | -1.1231493 | 2.57E-06 | 0.00052991 |
| ENSG00000196954 | CASP4 | -1.2257868 | 2.84E-06 | 0.00057716 |
| ENSG00000181381 | DDX60L | -1.2359637 | 2.96E-06 | 0.00059394 |
| ENSG00000171282 | RP11-1055B8.7 | -2.5766687 | 3.87E-06 | 0.0007644 |
| ENSG00000145990 | GFOD1 | -1.0346425 | 4.31E-06 | 0.00084024 |
| ENSG00000138642 | HERC6 | -1.4210999 | 4.50E-06 | 0.00086663 |
| ENSG00000145526 | CDH18 | -2.6931814 | 4.62E-06 | 0.00087846 |
| ENSG00000151882 | CCL28 | -1.4957715 | 4.74E-06 | 0.00088978 |
| ENSG00000231389 | HLA-DPA1 | -2.539317 | 5.78E-06 | 0.00104556 |
| ENSG00000138496 | PARP9 | -0.9470767 | 5.76E-06 | 0.00104556 |
| ENSG00000184979 | USP18 | -1.2927034 | 5.93E-06 | 0.00106023 |
| ENSG00000117228 | GBP1 | -1.8521315 | 6.04E-06 | 0.00106742 |
| ENSG00000165949 | IFI27 | -1.0566687 | 6.36E-06 | 0.00109628 |
| ENSG00000067066 | SP100 | -0.9837894 | 6.58E-06 | 0.00112114 |
| ENSG00000128342 | LIF | -2.195385 | 7.28E-06 | 0.00121305 |
| ENSG00000133106 | EPSTI1 | -1.8546361 | 7.52E-06 | 0.00122626 |
| ENSG00000135899 | SP110 | -1.0339483 | 7.47E-06 | 0.00122626 |
| ENSG00000169245 | CXCL10 | -2.6418469 | 7.62E-06 | 0.00122897 |
| ENSG00000138035 | PNPT1 | -0.8667033 | 8.17E-06 | 0.0012836 |
| ENSG00000138356 | AOX1 | -1.2972826 | 8.58E-06 | 0.00132585 |
| ENSG00000150594 | ADRA2A | -2.7714381 | 1.21E-05 | 0.00183079 |
| ENSG00000163131 | CTSS | -1.65193 | 1.31E-05 | 0.00194629 |
| ENSG00000171132 | PRKCE | -1.5534207 | 1.32E-05 | 0.00194629 |
| ENSG00000171159 | C9orf16 | -0.9108523 | 1.33E-05 | 0.00194629 |
| ENSG00000168394 | TAP1 | -1.2935997 | 1.43E-05 | 0.00208024 |
| ENSG00000100605 | ITPK1 | -0.9083033 | 1.57E-05 | 0.00225828 |
| ENSG00000172936 | MYD88 | -0.8339003 | 1.88E-05 | 0.00267946 |
| ENSG00000173559 | NABP1 | -1.8938713 | 1.96E-05 | 0.0027647 |
| ENSG00000213699 | SLC35F6 | -1.3917432 | 2.22E-05 | 0.00307607 |
| ENSG00000156587 | UBE2L6 | -1.1820935 | 2.24E-05 | 0.00307628 |
| ENSG00000178573 | MAF | -1.9559787 | 2.38E-05 | 0.00319868 |
| ENSG00000240184 | PCDHGC3 | -1.5311171 | 2.39E-05 | 0.00319868 |
| ENSG00000197415 | VEPH1 | -1.4882698 | 2.50E-05 | 0.00331615 |
| ENSG00000078081 | LAMP3 | -1.7533481 | 2.59E-05 | 0.00339698 |
| ENSG00000223808 | RP11-428L9.2 | -2.1760971 | 2.74E-05 | 0.00344907 |
| ENSG00000141449 | GREB1L | -1.5297249 | 2.69E-05 | 0.00344907 |
| ENSG00000023445 | BIRC3 | -0.9946572 | 2.72E-05 | 0.00344907 |
| ENSG00000160712 | IL6R | -1.3743498 | 2.83E-05 | 0.00352287 |
| ENSG00000010030 | ETV7 | -1.8387235 | 2.95E-05 | 0.00364179 |
| ENSG00000007944 | MYLIP | -1.1670875 | 3.03E-05 | 0.00368158 |
| ENSG00000170345 | FOS | -1.29656 | 3.15E-05 | 0.00380221 |
| ENSG00000177294 | FBXO39 | -2.7587208 | 3.28E-05 | 0.00386608 |
| ENSG00000132256 | TRIM5 | -1.080316 | 3.27E-05 | 0.00386608 |
| ENSG00000123609 | NMI | -1.0149481 | 3.25E-05 | 0.00386608 |
| ENSG00000043039 | BARX2 | -2.6364867 | 4.00E-05 | 0.00460098 |
| ENSG00000187017 | ESPN | -0.9841938 | 4.18E-05 | 0.00477459 |
| ENSG00000204592 | HLA-E | -1.5253427 | 4.24E-05 | 0.00479874 |
| ENSG00000137198 | GMPR | -1.6076953 | 4.43E-05 | 0.0048686 |
| ENSG00000111331 | OAS3 | -1.3602024 | 4.41E-05 | 0.0048686 |
| ENSG00000205629 | LCMT1 | -0.8237427 | 4.68E-05 | 0.00510091 |
| ENSG00000132821 | VSTM2L | -1.6966446 | 4.80E-05 | 0.00512091 |
| ENSG00000130589 | HELZ2 | -1.2220713 | 4.78E-05 | 0.00512091 |
| ENSG00000131242 | RAB11FIP4 | -0.8691401 | 4.79E-05 | 0.00512091 |
| ENSG00000143466 | IKBKE | -1.0730854 | 4.92E-05 | 0.00521289 |
| ENSG00000198624 | CCDC69 | -1.6361776 | 5.13E-05 | 0.00536289 |
| ENSG00000234745 | HLA-B | -1.6121484 | 5.11E-05 | 0.00536289 |
| ENSG00000068079 | IFI35 | -1.4525623 | 5.75E-05 | 0.00588569 |

#### TABLE S1 (continued)

| | | | | |
|---|---|---|---|---|
| ENSG00000128342 | LIF | -2.195385 | 7.28E-06 | 0.00121305 |
| ENSG00000133106 | EPSTI1 | -1.8546361 | 7.52E-06 | 0.00122626 |
| ENSG00000135899 | SP110 | -1.0339483 | 7.47E-06 | 0.00122626 |
| ENSG00000169245 | CXCL10 | -2.6418469 | 7.62E-06 | 0.00122897 |
| ENSG00000138035 | PNPT1 | -0.8667033 | 8.17E-06 | 0.0012836 |
| ENSG00000138356 | AOX1 | -1.2972826 | 8.58E-06 | 0.00132585 |
| ENSG00000150594 | ADRA2A | -2.7714381 | 1.21E-05 | 0.00183079 |
| ENSG00000163131 | CTSS | -1.65193 | 1.31E-05 | 0.00194629 |
| ENSG00000171132 | PRKCE | -1.5534207 | 1.32E-05 | 0.00194629 |
| ENSG00000171159 | C9orf16 | -0.9108523 | 1.33E-05 | 0.00194629 |
| ENSG00000168394 | TAP1 | -1.2935997 | 1.43E-05 | 0.00208024 |
| ENSG00000100605 | ITPK1 | -0.9083033 | 1.57E-05 | 0.00225828 |
| ENSG00000172936 | MYD88 | -0.8339003 | 1.88E-05 | 0.00267946 |
| ENSG00000173559 | NABP1 | -1.8938713 | 1.96E-05 | 0.0027647 |
| ENSG00000213699 | SLC35F6 | -1.3917432 | 2.22E-05 | 0.00307607 |
| ENSG00000156587 | UBE2L6 | -1.1820935 | 2.24E-05 | 0.00307628 |
| ENSG00000178573 | MAF | -1.9559787 | 2.38E-05 | 0.00319868 |
| ENSG00000240184 | PCDHGC3 | -1.5311171 | 2.39E-05 | 0.00319868 |
| ENSG00000197415 | VEPH1 | -1.4882698 | 2.50E-05 | 0.00331615 |
| ENSG00000078081 | LAMP3 | -1.7533481 | 2.59E-05 | 0.00339698 |
| ENSG00000223808 | RP11-428L9.2 | -2.1760971 | 2.74E-05 | 0.00344907 |
| ENSG00000141449 | GREB1L | -1.5297249 | 2.69E-05 | 0.00344907 |
| ENSG00000023445 | BIRC3 | -0.9946572 | 2.72E-05 | 0.00344907 |
| ENSG00000160712 | IL6R | -1.3743498 | 2.83E-05 | 0.00352287 |
| ENSG00000010030 | ETV7 | -1.8387235 | 2.95E-05 | 0.00364179 |
| ENSG00000007944 | MYLIP | -1.1670875 | 3.03E-05 | 0.00368158 |
| ENSG00000170345 | FOS | -1.29656 | 3.15E-05 | 0.00380221 |
| ENSG00000177294 | FBXO39 | -2.7587208 | 3.28E-05 | 0.00386608 |
| ENSG00000132256 | TRIM5 | -1.080316 | 3.27E-05 | 0.00386608 |
| ENSG00000123609 | NMI | -1.0149481 | 3.25E-05 | 0.00386608 |
| ENSG00000043039 | BARX2 | -2.6364867 | 4.00E-05 | 0.00460098 |
| ENSG00000187017 | ESPN | -0.9841938 | 4.18E-05 | 0.00477459 |
| ENSG00000204592 | HLA-E | -1.5253427 | 4.24E-05 | 0.00479874 |
| ENSG00000137198 | GMPR | -1.6076953 | 4.43E-05 | 0.0048686 |
| ENSG00000111331 | OAS3 | -1.3602024 | 4.41E-05 | 0.0048686 |
| ENSG00000205629 | LCMT1 | -0.8237427 | 4.68E-05 | 0.00510091 |
| ENSG00000132821 | VSTM2L | -1.6966446 | 4.80E-05 | 0.00512091 |
| ENSG00000130589 | HELZ2 | -1.2220713 | 4.78E-05 | 0.00512091 |
| ENSG00000131242 | RAB11FIP4 | -0.8691401 | 4.79E-05 | 0.00512091 |
| ENSG00000143466 | IKBKE | -1.0730854 | 4.92E-05 | 0.00521289 |
| ENSG00000198624 | CCDC69 | -1.6361776 | 5.13E-05 | 0.00536289 |
| ENSG00000234745 | HLA-B | -1.6121484 | 5.11E-05 | 0.00536289 |
| ENSG00000068079 | IFI35 | -1.4525623 | 5.75E-05 | 0.00588569 |
| ENSG00000019582 | CD74 | -2.9975606 | 6.04E-05 | 0.0061371 |
| ENSG00000163814 | CDCP1 | -0.7662169 | 6.08E-05 | 0.0061371 |
| ENSG00000029153 | ARNTL2 | -3.0010535 | 6.16E-05 | 0.00617756 |
| ENSG00000104142 | VPS18 | -0.820787 | 6.25E-05 | 0.00622407 |
| ENSG00000185404 | SP140L | -0.9738197 | 6.56E-05 | 0.00648369 |
| ENSG00000036672 | USP2 | -1.8969455 | 6.93E-05 | 0.00679168 |
| ENSG00000174808 | BTC | -1.7189536 | 6.96E-05 | 0.00679168 |
| ENSG00000173821 | RNF213 | -0.760724 | 7.17E-05 | 0.00694986 |
| ENSG00000137462 | TLR2 | -1.8509398 | 7.38E-05 | 0.00707012 |
| ENSG00000157557 | ETS2 | -1.065431 | 7.72E-05 | 0.00733619 |
| ENSG00000164756 | SLC30A8 | -2.9013221 | 7.77E-05 | 0.00734085 |
| ENSG00000154478 | GPR26 | -3.4540666 | 8.40E-05 | 0.00783627 |
| ENSG00000168539 | CHRM1 | -1.7183495 | 8.57E-05 | 0.00789321 |
| ENSG00000110057 | UNC93B1 | -0.8867719 | 8.53E-05 | 0.00789321 |
| ENSG00000204642 | HLA-F | -2.0074191 | 8.69E-05 | 0.00790569 |
| ENSG00000137965 | IFI44 | -1.7818806 | 8.67E-05 | 0.00790569 |
| ENSG00000145476 | CYP4V2 | -1.6865153 | 9.01E-05 | 0.00808679 |
| ENSG00000165194 | PCDH19 | -1.4916956 | 8.99E-05 | 0.00808679 |
| ENSG00000067082 | KLF6 | -1.0997662 | 9.05E-05 | 0.00808679 |

#### TABLE S1 (continued)

| | | | | |
|---|---|---|---|---|
| ENSG00000100342 | APOL1 | -1.5223418 | 9.11E-05 | 0.00808892 |
| ENSG00000204257 | HLA-DMA | -2.3561673 | 9.44E-05 | 0.00828158 |
| ENSG00000162654 | GBP4 | -1.4858828 | 9.78E-05 | 0.00853445 |
| ENSG00000108679 | LGALS3BP | -0.8408733 | 0.00010922 | 0.00947434 |
| ENSG00000184545 | DUSP8 | -1.3330032 | 0.00011373 | 0.00980797 |
| ENSG00000043143 | JADE2 | -0.6889421 | 0.00011815 | 0.01013045 |
| ENSG00000120129 | DUSP1 | -1.3390559 | 0.00013596 | 0.01159056 |
| ENSG00000128274 | A4GALT | -1.5029957 | 0.00014277 | 0.01210105 |
| ENSG00000138074 | SLC5A6 | -0.9047347 | 0.00015024 | 0.01266235 |
| ENSG00000197102 | DYNC1H1 | -0.750662 | 0.00015218 | 0.01268149 |
| ENSG00000164713 | BRI3 | -0.9715575 | 0.00015414 | 0.01277273 |
| ENSG00000244588 | RAD21L1 | -1.9014702 | 0.00015671 | 0.01291409 |
| ENSG00000172458 | IL17D | -1.741006 | 0.00016391 | 0.01333717 |
| ENSG00000107201 | DDX58 | -1.0848571 | 0.00016455 | 0.01333717 |
| ENSG00000166165 | CKB | -0.8487911 | 0.00016547 | 0.01333912 |
| ENSG00000134326 | CMPK2 | -3.5010975 | 0.00016801 | 0.01347043 |
| ENSG00000124201 | ZNFX1 | -0.8983295 | 0.00017042 | 0.0135908 |
| ENSG00000184602 | SNN | -0.9246791 | 0.00017367 | 0.01377537 |
| ENSG00000111799 | COL12A1 | -1.1187756 | 0.00017488 | 0.01379796 |
| ENSG00000107731 | UNC5B | -1.2114802 | 0.00017695 | 0.01388724 |
| ENSG00000174718 | KIAA1551 | -0.9690174 | 0.00017929 | 0.01392368 |
| ENSG00000123094 | RASSF8 | -1.1839099 | 0.00018129 | 0.01400562 |
| ENSG00000143369 | ECM1 | -1.6433724 | 0.00018798 | 0.01441572 |
| ENSG00000117266 | CDK18 | -1.0437284 | 0.00018854 | 0.01441572 |
| ENSG00000073464 | CLCN4 | -2.3534639 | 0.00019016 | 0.01446473 |
| ENSG00000102032 | RENBP | -1.9728087 | 0.00019736 | 0.01485998 |
| ENSG00000128335 | APOL2 | -1.3397838 | 0.00020074 | 0.01503828 |
| ENSG00000107798 | LIPA | -0.753269 | 0.00021252 | 0.01576173 |
| ENSG00000163235 | TGFA | -0.9704612 | 0.00022909 | 0.01685023 |
| ENSG00000197355 | UAP1L1 | -0.9552212 | 0.00022947 | 0.01685023 |
| ENSG00000129187 | DCTD | -0.8482121 | 0.00023613 | 0.01725388 |
| ENSG00000168143 | FAM83B | -1.0638682 | 0.0002414 | 0.017467 |
| ENSG00000130522 | JUND | -0.8241667 | 0.00025639 | 0.01846162 |
| ENSG00000100911 | PSME2 | -0.795382 | 0.00026401 | 0.01882716 |
| ENSG00000126709 | IFI6 | -1.2600284 | 0.0002681 | 0.01893706 |
| ENSG00000132109 | TRIM21 | -1.2476931 | 0.000267 | 0.01893706 |
| ENSG00000170581 | STAT2 | -0.8166153 | 0.00027301 | 0.01901215 |
| ENSG00000107537 | PHYH | -1.4474664 | 0.00029987 | 0.02068827 |
| ENSG00000163840 | DTX3L | -0.7681021 | 0.00030182 | 0.0207139 |
| ENSG00000162889 | MAPKAPK2 | -0.695376 | 0.00030303 | 0.0207139 |
| ENSG00000121064 | SCPEP1 | -1.1547825 | 0.00030456 | 0.02072293 |
| ENSG00000182952 | HMGN4 | -0.9792485 | 0.00033285 | 0.02244153 |
| ENSG00000152894 | PTPRK | -0.7683928 | 0.00033457 | 0.02245522 |
| ENSG00000183908 | LRRC55 | -2.6545779 | 0.00033822 | 0.02249669 |
| ENSG00000072110 | ACTN1 | -0.7293092 | 0.00033724 | 0.02249669 |
| ENSG00000177628 | GBA | -0.7686344 | 0.00035188 | 0.02330105 |
| ENSG00000003436 | TFPI | -2.7231521 | 0.00035355 | 0.02330763 |
| ENSG00000100292 | HMOX1 | -1.6060696 | 0.00036923 | 0.02392582 |
| ENSG00000169499 | PLEKHA2 | -0.8950884 | 0.00036689 | 0.02392582 |
| ENSG00000132481 | TRIM47 | -0.839653 | 0.00037799 | 0.0243769 |
| ENSG00000013619 | MAMLD1 | -2.0487447 | 0.00038528 | 0.02473986 |
| ENSG00000151322 | NPAS3 | -1.9307766 | 0.0004128 | 0.02616703 |
| ENSG00000205336 | ADGRG1 | -0.6735176 | 0.00041148 | 0.02616703 |
| ENSG00000125775 | SDCBP2 | -1.3318725 | 0.00041798 | 0.02638269 |
| ENSG00000119655 | NPC2 | -0.8546411 | 0.00043332 | 0.02712029 |
| ENSG00000103249 | CLCN7 | -0.8728197 | 0.0004671 | 0.02898954 |
| ENSG00000135048 | TMEM2 | -1.2970391 | 0.00047408 | 0.02907899 |
| ENSG00000130813 | C19orf66 | -1.0906674 | 0.00047169 | 0.02907899 |
| ENSG00000013364 | MVP | -0.6785436 | 0.00047442 | 0.02907899 |
| ENSG00000179431 | FJX1 | -1.1240836 | 0.00048926 | 0.02986506 |

### TABLE S1 (continued)

| | | | | |
|---|---|---|---|---|
| ENSG00000135127 | BICDL1 | -1.2243237 | 0.00051421 | 0.03113168 |
| ENSG00000166710 | B2M | -1.0155833 | 0.00052758 | 0.03181151 |
| ENSG00000197081 | IGF2R | -0.7424498 | 0.00053028 | 0.03184478 |
| ENSG00000184216 | IRAK1 | -0.6693657 | 0.00054268 | 0.03236765 |
| ENSG00000185896 | LAMP1 | -0.6377334 | 0.00054553 | 0.03236765 |
| ENSG00000130766 | SESN2 | -1.317198 | 0.00055244 | 0.03264654 |
| ENSG00000164342 | TLR3 | -1.2913436 | 0.00055783 | 0.03270498 |
| ENSG00000162976 | PQLC3 | -1.4171628 | 0.00058662 | 0.03323839 |
| ENSG00000163297 | ANTXR2 | -1.4106415 | 0.00057807 | 0.03323839 |
| ENSG00000173221 | GLRX | -0.9651887 | 0.00057888 | 0.03323839 |
| ENSG00000068650 | ATP11A | -0.8477554 | 0.00058514 | 0.03323839 |
| ENSG00000065923 | SLC9A7 | -0.6769479 | 0.00057598 | 0.03323839 |
| ENSG00000168003 | SLC3A2 | -1.1150234 | 0.000595 | 0.03355749 |
| ENSG00000164506 | STXBP5 | -1.045298 | 0.00060553 | 0.03365217 |
| ENSG00000182022 | CHST15 | -0.8626274 | 0.00060575 | 0.03365217 |
| ENSG00000166900 | STX3 | -0.7712919 | 0.00060048 | 0.03365217 |
| ENSG00000177556 | ATOX1 | -0.7089019 | 0.00061162 | 0.03385121 |
| ENSG00000185885 | IFITM1 | -1.0897119 | 0.00061696 | 0.03401969 |
| ENSG00000128578 | STRIP2 | -1.5751092 | 0.00064479 | 0.03529187 |
| ENSG00000204131 | NHSL2 | -1.7299957 | 0.00065727 | 0.03584279 |
| ENSG00000213694 | S1PR3 | -0.776145 | 0.00065982 | 0.03584998 |
| ENSG00000164292 | RHOBTB3 | -0.8431656 | 0.00067957 | 0.03637869 |
| ENSG00000176463 | SLCO3A1 | -1.5525233 | 0.00069262 | 0.0364311 |
| ENSG00000177606 | JUN | -1.4303474 | 0.00069007 | 0.0364311 |
| ENSG00000082482 | KCNK2 | -1.1879226 | 0.00068682 | 0.0364311 |
| ENSG00000027697 | IFNGR1 | -0.7758301 | 0.00069778 | 0.03657322 |
| ENSG00000160932 | LY6E | -0.7804205 | 0.00070849 | 0.03700344 |
| ENSG00000012223 | LTF | -2.4492247 | 0.00072824 | 0.03790172 |
| ENSG00000139112 | GABARAPL1 | -2.0369819 | 0.00073523 | 0.03808251 |
| ENSG00000142089 | IFITM3 | -1.0637565 | 0.00073685 | 0.03808251 |
| ENSG00000169851 | PCDH7 | -3.7670061 | 0.00074365 | 0.03830032 |
| ENSG00000008735 | MAPK8IP2 | -1.4246212 | 0.00077215 | 0.0393584 |
| ENSG00000213689 | TREX1 | -1.1940332 | 0.00077099 | 0.0393584 |
| ENSG00000124275 | MTRR | -0.6260982 | 0.00078931 | 0.03982282 |
| ENSG00000101474 | APMAP | -0.6504945 | 0.00079425 | 0.03993617 |
| ENSG00000159461 | AMFR | -0.8176736 | 0.0008145 | 0.04067832 |
| ENSG00000184557 | SOCS3 | -1.476266 | 0.00082747 | 0.04118764 |
| ENSG00000128284 | APOL3 | -1.3470228 | 0.00083549 | 0.04144752 |
| ENSG00000116729 | WLS | -1.5318873 | 0.00084911 | 0.04184326 |
| ENSG00000089327 | FXYD5 | -0.8589102 | 0.00087511 | 0.04242016 |
| ENSG00000105711 | SCN1B | -1.2840985 | 0.00088737 | 0.04265319 |
| ENSG00000137714 | FDX1 | -1.3598626 | 0.00090558 | 0.04333075 |
| ENSG00000223573 | TINCR | -0.8142372 | 0.00091912 | 0.04383716 |
| ENSG00000136717 | BIN1 | -1.011929 | 0.00094047 | 0.04456843 |
| ENSG00000150556 | LYPD6B | -1.1381476 | 0.00097261 | 0.04579903 |
| ENSG00000167552 | TUBA1A | -1.3204515 | 0.00099318 | 0.04647245 |
| ENSG00000122643 | NT5C3A | -0.8205239 | 0.00104893 | 0.04848374 |
| ENSG00000182179 | UBA7 | -0.9532607 | 0.00106574 | 0.04897244 |
| ENSG00000198431 | TXNRD1 | -1.0826951 | 0.00109505 | 0.04997787 |

### TABLE S2

## Up in 68H (84)

| ensembl_gene_id | gene_symbol | log2FoldChange | pvalue | padj |
|---|---|---|---|---|
| ENSG00000068078 | FGFR3 | 1.48987281 | 8.26E-12 | 1.53E-08 |
| ENSG00000100033 | PRODH | 2.01442889 | 9.21E-10 | 9.11E-07 |
| ENSG00000149328 | GLB1L2 | 2.83339121 | 6.27E-08 | 2.66E-05 |
| ENSG00000130427 | EPO | 5.19598155 | 7.88E-08 | 3.16E-05 |
| ENSG00000160862 | AZGP1 | 1.49574867 | 1.47E-07 | 4.97E-05 |
| ENSG00000170786 | SDR16C5 | 1.6721625 | 2.88E-07 | 9.28E-05 |
| ENSG00000126562 | WNK4 | 2.34556711 | 3.88E-07 | 0.00011301 |
| ENSG00000230937 | MIR205HG | 1.51932697 | 5.39E-07 | 0.0001427 |
| ENSG00000147234 | FRMPD3 | 1.74275563 | 5.97E-07 | 0.00015548 |
| ENSG00000112655 | PTK7 | 1.34007752 | 8.61E-07 | 0.00021652 |
| ENSG00000056998 | GYG2 | 2.05866682 | 1.16E-06 | 0.0002867 |
| ENSG00000131620 | ANO1 | 1.97823923 | 1.39E-06 | 0.00033737 |
| ENSG00000165125 | TRPV6 | 1.87249441 | 1.55E-06 | 0.00036118 |
| ENSG00000105048 | TNNT1 | 1.3492053 | 1.60E-06 | 0.0003652 |
| ENSG00000178538 | CA8 | 1.87168089 | 1.66E-06 | 0.0003728 |
| ENSG00000147852 | VLDLR | 2.81827891 | 5.76E-06 | 0.00104556 |
| ENSG00000159763 | PIP | 1.90510229 | 6.17E-06 | 0.00107683 |
| ENSG00000117154 | IGSF21 | 4.08939172 | 6.69E-06 | 0.00112706 |
| ENSG00000166106 | ADAMTS15 | 1.65891446 | 7.78E-06 | 0.00124042 |
| ENSG00000124126 | PREX1 | 1.37682 | 8.22E-06 | 0.0012836 |
| ENSG00000134323 | MYCN | 1.54267928 | 9.45E-06 | 0.00144543 |
| ENSG00000010310 | GIPR | 1.10396821 | 2.19E-05 | 0.00306914 |
| ENSG00000110484 | SCGB2A2 | 2.92405722 | 2.33E-05 | 0.00317251 |
| ENSG00000147257 | GPC3 | 2.37283515 | 2.63E-05 | 0.00342004 |
| ENSG00000172828 | CES3 | 2.86187452 | 2.73E-05 | 0.00344907 |
| ENSG00000179820 | MYADM | 1.19957828 | 3.00E-05 | 0.00367686 |
| ENSG00000136883 | KIF12 | 1.50566495 | 3.76E-05 | 0.0043963 |
| ENSG00000197520 | FAM177B | 4.03516418 | 3.90E-05 | 0.00451871 |
| ENSG00000139865 | TTC6 | 0.99926941 | 4.32E-05 | 0.00483446 |
| ENSG00000196482 | ESRRG | 1.11138932 | 4.33E-05 | 0.00483446 |
| ENSG00000108947 | EFNB3 | 1.9049157 | 5.58E-05 | 0.00578409 |
| ENSG00000152256 | PDK1 | 1.55267503 | 5.63E-05 | 0.00579985 |
| ENSG00000166927 | MS4A7 | 1.61002918 | 7.39E-05 | 0.00707012 |
| ENSG00000139631 | CSAD | 0.81863133 | 8.22E-05 | 0.00771918 |
| ENSG00000149403 | GRIK4 | 1.65681162 | 9.28E-05 | 0.00818956 |
| ENSG00000230551 | CTB-89H12.4 | 0.68685207 | 0.00015137 | 0.01268149 |
| ENSG00000203668 | CHML | 0.76991079 | 0.00015804 | 0.01295121 |
| ENSG00000196074 | SYCP2 | 0.83822177 | 0.00017815 | 0.01390822 |
| ENSG00000236467 | KCNMA1-AS1 | 1.86097473 | 0.00019294 | 0.01460115 |
| ENSG00000260711 | RP11-747H7.3 | 0.8259621 | 0.0002125 | 0.01576173 |
| ENSG00000129682 | FGF13 | 1.7621803 | 0.00023844 | 0.0173368 |
| ENSG00000050438 | SLC4A8 | 0.90639354 | 0.000259 | 0.01855888 |
| ENSG00000183888 | C1orf64 | 1.71099089 | 0.00027085 | 0.01895077 |
| ENSG00000158747 | NBL1 | 1.84768242 | 0.00028902 | 0.02003278 |
| ENSG00000082397 | EPB41L3 | 1.53285418 | 0.00031373 | 0.0212493 |
| ENSG00000266714 | MYO15B | 0.74532279 | 0.0003579 | 0.02348974 |
| ENSG00000087495 | PHACTR3 | 2.33565584 | 0.00036938 | 0.02392582 |
| ENSG00000008513 | ST3GAL1 | 1.37465217 | 0.00041262 | 0.02616703 |
| ENSG00000250397 | RP11-1391J7.1 | 1.3306612 | 0.0004319 | 0.02712029 |
| ENSG00000248771 | LINC01207 | 2.2457356 | 0.00046079 | 0.02871781 |
| ENSG00000213700 | RPL17P50 | 2.0811808 | 0.00051013 | 0.03101144 |
| ENSG00000102001 | CACNA1F | 1.43851869 | 0.00054435 | 0.03236765 |
| ENSG00000259964 | THSD4-AS1 | 1.31373927 | 0.00055686 | 0.03270498 |
| ENSG00000182257 | PRR34 | 1.06410572 | 0.0005751 | 0.03323839 |
| ENSG00000100234 | TIMP3 | 1.47267282 | 0.00058153 | 0.03323839 |

#### TABLE S2 (continued)

| | | | | |
|---|---|---|---|---|
| ENSG00000189058 | APOD | 1.58920405 | 0.0005871 | 0.03323839 |
| ENSG00000131969 | ABHD12B | 1.72657595 | 0.00056946 | 0.03323839 |
| ENSG00000135052 | GOLM1 | 1.21597689 | 0.00060124 | 0.03365217 |
| ENSG00000198363 | ASPH | 1.37407637 | 0.00062629 | 0.03440653 |
| ENSG00000177685 | CRACR2B | 0.84826759 | 0.00067171 | 0.03636319 |
| ENSG00000170476 | MZB1 | 0.9587559 | 0.00068181 | 0.03637869 |
| ENSG00000164199 | ADGRV1 | 1.24495619 | 0.00067455 | 0.03637869 |
| ENSG00000229028 | KRT223P | 2.22416102 | 0.00067829 | 0.03637869 |
| ENSG00000104267 | CA2 | 2.95359698 | 0.00069096 | 0.0364311 |
| ENSG00000075891 | PAX2 | 2.66937856 | 0.00076734 | 0.0393584 |
| ENSG00000164742 | ADCY1 | 1.56946363 | 0.00077733 | 0.03948684 |
| ENSG00000148488 | ST8SIA6 | 0.9906507 | 0.00078242 | 0.03960948 |
| ENSG00000188206 | HNRNPU-AS1 | 1.01669863 | 0.00080421 | 0.04029998 |
| ENSG00000166068 | SPRED1 | 1.84528663 | 0.00083866 | 0.04146597 |
| ENSG00000162482 | AKR7A3 | 1.66056782 | 0.00085512 | 0.04190822 |
| ENSG00000101825 | MXRA5 | 2.81137494 | 0.00085608 | 0.04190822 |
| ENSG00000188365 | AC092171.2 | 0.88914641 | 0.0008736 | 0.04242016 |
| ENSG00000254440 | PBOV1 | 1.34327992 | 0.00086955 | 0.04242016 |
| ENSG00000099954 | CECR2 | 1.04254687 | 0.00088789 | 0.04265319 |
| ENSG00000170525 | PFKFB3 | 1.23831566 | 0.00088855 | 0.04265319 |
| ENSG00000005379 | TSPOAP1 | 0.75579225 | 0.00092529 | 0.04398981 |
| ENSG00000037965 | HOXC8 | 0.75512249 | 0.00095114 | 0.04493073 |
| ENSG00000144218 | AFF3 | 1.34927717 | 0.00098383 | 0.04618094 |
| ENSG00000253154 | CTA-392E5.1 | 3.56856017 | 0.00101293 | 0.04724775 |
| ENSG00000131746 | TNS4 | 1.18404645 | 0.00102931 | 0.04786139 |
| ENSG00000141753 | IGFBP4 | 0.60688871 | 0.00104923 | 0.04848374 |
| ENSG00000117983 | MUC5B | 1.00185793 | 0.00106641 | 0.04897244 |
| ENSG00000123572 | NRK | 1.46099236 | 0.0010795 | 0.04942045 |

#### TABLE S3

GSEA analysis with genes downregulated in 68H compared to 68N

| Gene Set Name | # Genes in Gene Set (K) | # Genes in Overlap (k) | k/K | p-value | FDR q-value |
|---|---|---|---|---|---|
| HALLMARK_INTERFRON_GAMMA_RESPONSE | 200 | 50 | 0.25 | 2.91E-71 | 2.47E-68 |
| HALLMARK_INTERFERON_ALPHA_RESPONSE | 97 | 39 | 0.40 | 4.74E-65 | 2.02E-62 |
| STTTCRNTTT_IRF_Q6 | 188 | 20 | 0.106 | 6.17E-21 | 1.75E-18 |
| ISRE_01 | 247 | 21 | 0.085 | 7.11E-20 | 1.51E-17 |
| IRF7_01 | 252 | 21 | 0.083 | 1.08E-19 | 1.84E-17 |

#### TABLE S4

GSEA analysis with genes upregulated in 68H compared to 68N

| Gene Set Name | # Genes in Gene Set (K) | # Genes in Overlap (k) | k/K | p-value | FDR q-value |
|---|---|---|---|---|---|
| HALLMARK_KRAS_SIGNALING_UP | 200 | 5 | 0.025 | 1.15E-05 | 2.72E-03 |
| HALLMARK_ESTROGEN_RESPONSE_LATE | 200 | 4 | 0.020 | 2.15E-04 | 1.69E-02 |
| HALLMARK_HYPOXIA | 200 | 4 | 0.020 | 2.15E-04 | 1.69E-02 |
| ELVIDGE_HYPOXIA_UP | 172 | 4 | 0.023 | 4.49E-04 | 3.68E-02 |

### TABLE S5

GSEA gene sets showing significant overlap with genes overexpressed in responders (top 20 shown here)

| Gene Set Name | # Genes in Gene Set (K) | Description | # Genes in Overlap (k) | k/K | p-value | FDR q-value |
|---|---|---|---|---|---|---|
| **CHARAFE_BREAST_CANCER_LUMINAL_VS_MESENCHYMAL_UP** | **451** | **Genes up-regulated in luminal-like breast cancer cell lines compared to the mesenchymal-like ones.** | **118** | **0.2616** | **2.53E-136** | **2.39E-132** |
| **HOLLERN_EMT_BREAST_TUMOR_DN** | **118** | **Genes that that have low expression in mammary tumors of epithelial-mesenchymal transition (EMT) histology.** | **63** | **0.5339** | **1.65E-95** | **7.78E-92** |
| COLDREN_GEFITINIB_RESISTANCE_DN | 225 | Genes down-regulated in NSCLC (non-small cell lung carcinoma) cell lines resistant to gefitinib [PubChem=123631] compared to the sensitive ones. | 66 | 0.2933 | 1.65E-78 | 5.18E-75 |
| **ONDER_CDH1_TARGETS_2_DN** | **473** | **Genes down-regulated in HMLE cells (immortalized nontransformed mammary epithelium) after E-cadhedrin (CDH1) [GeneID=999] knockdown by RNAi.** | **76** | **0.1607** | **6.31E-69** | **1.49E-65** |
| CHARAFE_BREAST_CANCER_BASAL_VS_MESENCHYMAL_UP | 125 | Genes up-regulated in basal-like breast cancer cell lines as compared to the mesenchymal-like ones. | 45 | 0.36 | 3.31E-58 | 6.25E-55 |
| LIM_MAMMARY_STEM_CELL_DN | 408 | Genes consistently down-regulated in mammary stem cells both in mouse and human species. | 63 | 0.1544 | 8.03E-56 | 1.26E-52 |
| LEE_BMP2_TARGETS_UP | 752 | Genes up-regulated in uterus upon knockout of BMP2 [GeneID=650]. | 69 | 0.0918 | 1.12E-45 | 1.51E-42 |
| MEISSNER_BRAIN_HCP_WITH_H3K4ME3_AND_H3K27ME3 | 1056 | Genes with high-CpG-density promoters (HCP) bearing histone H3 dimethylation at K4 (H3K4me2) and trimethylation at K27 (H3K27me3) in brain. | 76 | 0.072 | 7.48E-43 | 8.82E-40 |
| JAEGER_METASTASIS_DN | 258 | Genes down-regulated in metastases from malignant melanoma compared to the primary tumors. | 43 | 0.1667 | 1.27E-39 | 1.33E-36 |
| DODD_NASOPHARYNGEAL_CARCINOMA_UP | 1785 | Genes up-regulated in nasopharyngeal carcinoma (NPC) compared to the normal tissue. | 87 | 0.0487 | 4.50E-36 | 4.25E-33 |
| MCBRYAN_PUBERTAL_BREAST_4_5WK_UP | 259 | Genes up-regulated during pubertal mammary gland development between week 4 and 5. | 39 | 0.1506 | 3.14E-34 | 2.69E-31 |
| **CHARAFE_BREAST_CANCER_LUMINAL_VS_BASAL_UP** | **383** | **Genes up-regulated in luminal-like breast cancer cell lines compared to the basal-like ones.** | **44** | **0.1149** | **2.49E-33** | **1.96E-30** |
| HALLMARK_ESTROGEN_RESPONSE_EARLY | 200 | Genes defining early response to estrogen. | 34 | 0.17 | 8.33E-32 | 6.04E-29 |
| **AIGNER_ZEB1_TARGETS** | **35** | **Genes up-regulated in MDA-MB-231 cells (breast cancer) after knockdown of ZEB1 [GeneID=6935] by RNAi.** | **19** | **0.5429** | **1.60E-29** | **1.08E-26** |
| MEISSNER_NPC_HCP_WITH_H3_UNMETHYLATED | 526 | Genes with high-CpG-density promoters (HCP) that have no histone H3 methylation marks in neural precursor cells (NPC). | 45 | 0.0856 | 1.63E-28 | 1.03E-25 |
| LIEN_BREAST_CARCINOMA_METAPLASTIC_VS_DUCTAL_DN | 112 | Genes down-regulated between two breast carcinoma subtypes: metaplastic (MCB) and ductal (DCB). | 26 | 0.2321 | 2.49E-28 | 1.47E-25 |
| GOZGIT_ESR1_TARGETS_DN | 767 | Genes down-regulated in TMX2-28 cells (breast cancer) which do not express ESR1 [GeneID=2099]) compared to the parental MCF7 cells which do. | 51 | 0.0665 | 4.46E-27 | 2.47E-24 |
| **SMID_BREAST_CANCER_BASAL_DN** | **699** | **Genes down-regulated in basal subtype of breast cancer samles.** | **46** | **0.0658** | **2.73E-24** | **1.43E-21** |
| CHEMNITZ_RESPONSE_TO_PROSTAGLANDIN_E2_DN | 389 | Genes down-regulated in CD4+ [GeneID=920] T lymphocytes after stimulation with prostaglandin E2 [PubChem=5280360]. | 36 | 0.0925 | 3.85E-24 | 1.91E-21 |
| WAMUNYOKOLI_OVARIAN_CANCER_LMP_UP | 270 | Genes up-regulated in mucinous ovarian carcinoma tumors of low malignant potential (LMP) compared to normal ovarian surface epithelium tissue. | 31 | 0.1148 | 9.61E-24 | 4.53E-21 |

### TABLE S6

GSEA gene sets showing significant overlap with overexpressed in non-responders (top 20 shown + hypoxia pathway shown here)

| Gene Set Name | # Genes in Gene Set (K) | Description | # Genes in Overlap (k) | k/K | p-value | FDR q-value |
|---|---|---|---|---|---|---|
| **CHARAFE_BREAST_CANCER_LUMINAL_VS_MESENCHYMAL_DN** | **465** | **Genes down-regulated in luminal-like breast cancer cell lines compared to the mesenchymal-like ones.** | **79** | **0.1699** | **1.83E-90** | **1.73E-86** |
| SCHUETZ_BREAST_CANCER_DUCTAL_INVASIVE_UP | 352 | Genes up-regulated in invasive ductal carcinoma (IDC) relative to ductal carcinoma in situ (DCIS, non-invasive). | 48 | 0.1364 | 2.64E-49 | 1.25E-45 |
| LIM_MAMMARY_STEM_CELL_UP | 470 | Genes consistently up-regulated in mammary stem cells both in mouse and human species. | 48 | 0.1021 | 3.71E-43 | 1.16E-39 |
| **HOLLERN_EMT_BREAST_TUMOR_UP** | **135** | **Genes that are highly expressed in mammary tumors of epithelial-mesenchymal transition (EMT) histology.** | **29** | **0.2148** | **5.23E-36** | **1.23E-32** |
| **ONDER_CDH1_TARGETS_2_UP** | **256** | **Genes up-regulated in HMLE cells (immortalized nontransformed mammary epithelium) after E-cadhedrin (CDH1) [GeneID=999] knockdown by RNAi.** | **34** | **0.1328** | **1.46E-34** | **2.75E-31** |
| **HALLMARK_EPITHELIAL_MESENCHYMAL_TRANSITION** | **200** | **Genes defining epithelial-mesenchymal transition, as in wound healing, fibrosis and metastasis.** | **31** | **0.155** | **9.87E-34** | **1.55E-30** |
| CHICAS_RB1_TARGETS_CONFLUENT | 563 | Genes up-regulated in confluent IMR90 cells (fibroblast) after knockdown of RB1 [GeneID=5925] by RNAi. | 41 | 0.0728 | 6.39E-31 | 8.61E-28 |
| LIU_PROSTATE_CANCER_DN | 491 | Genes down-regulated in prostate cancer samples. | 37 | 0.0754 | 1.81E-28 | 2.13E-25 |
| **CHARAFE_BREAST_CANCER_LUMINAL_VS_BASAL_DN** | **454** | **Genes down-regulated in luminal-like breast cancer cell lines compared to the basal-like ones.** | **35** | **0.0771** | **2.70E-27** | **2.83E-24** |
| LINDGREN_BLADDER_CANCER_CLUSTER_2B | 388 | Genes specifically up-regulated in Cluster IIb of urothelial cell carcinom (UCC) tumors. | 33 | 0.0851 | 3.82E-27 | 3.60E-24 |
| MEISSNER_BRAIN_HCP_WITH_H3K4ME3_AND_H3K27ME3 | 1056 | Genes with high-CpG-density promoters (HCP) bearing histone H3 dimethylation at K4 (H3K4me2) and trimethylation at K27 (H3K27me3) in brain. | 46 | 0.0436 | 4.49E-25 | 3.85E-22 |
| AACTTT_UNKNOWN | 1936 | Genes having at least one occurence of the highly conserved motif M17 AACTTT in the region spanning up to 4 kb around their transcription start sites. The motif does not match any known transcription factor binding site (v7.4 TRANSFAC). | 59 | 0.0305 | 3.19E-24 | 2.51E-21 |
| SMID_BREAST_CANCER_NORMAL_LIKE_UP | 482 | Genes up-regulated in the normal-like subtype of breast cancer. | 33 | 0.0685 | 3.91E-24 | 2.84E-21 |
| SMID_BREAST_CANCER_LUMINAL_B_DN | 586 | Genes down-regulated in the luminal B subtype of breast cancer. | 35 | 0.0597 | 1.37E-23 | 9.26E-21 |
| BOQUEST_STEM_CELL_UP | 262 | Genes up-regulated in freshly isolated CD31- [GeneID=5175] (stromal stem cells from adipose tissue) versus the CD31+ (non-stem) counterparts. | 25 | 0.0954 | 5.21E-22 | 3.28E-19 |
| TGGAAA_NFAT_Q4_01 | 1927 | Genes having at least one occurence of the highly conserved motif M55 TGGAAA sites. The motif matches transcription factor binding site V$NFAT_Q4_01 (v7.4 TRANSFAC). | 56 | 0.0291 | 5.59E-22 | 3.30E-19 |
| REN_ALVEOLAR_RHABDOMYOSARCOMA_DN | 407 | Genes commonly down-regulated in human alveolar rhabdomyosarcoma (ARMS) and its mouse model overexpressing PAX3-FOXO1 [GeneID=5077;2308] fusion. | 29 | 0.0713 | 8.66E-22 | 4.81E-19 |
| CHEN_METABOLIC_SYNDROM_NETWORK | 1213 | Genes forming the macrophage-enriched metabolic network (MEMN) claimed to have a causal relationship with the metabolic syndrom traits. | 45 | 0.0371 | 9.79E-22 | 5.13E-19 |
| OISHI_CHOLANGIOMA_STEM_CELL_LIKE_DN | 275 | Genes under-expressed in stem cell-like cholangiocellular carcinoma | 25 | 0.0909 | 1.72E-21 | 8.56E-19 |
| PASINI_SUZ12_TARGETS_DN | 309 | Genes down-regulated in ES (embryonic stem cells) with defficient SUZ12 [GeneID=23512]. | 26 | 0.0841 | 1.88E-21 | 8.88E-19 |
| **MANALO_HYPOXIA_UP** | 204 | Genes up-regulated in response to both hypoxia and overexpression of an active form of HIF1A [GeneID=3091]. | 17 | 0.0833 | 2.34E-14 | 4.70E-12 |

**TABLE S7**

Genes belonging to the hypoxia and IFN/AP signatures

#### Hypoxia

| | |
|---|---|
| VEGFA | C20orf20 |
| SLC2A1 | HIG2 |
| PGAM1 | GAPDH |
| ENO1 | MRPL13 |
| LDHA | CHCHD2 |
| TPI1 | YKT6 |
| P4HA1 | NP |
| MRPS17 | CORO1C |
| CDKN3 | SEC61G |
| ADM | ANKRD37 |
| NDRG1 | ESRP1 |
| TUBB6 | PFKP |
| ALDOA | SHCBP1 |
| MIF | CTSL2 |
| ACOT7 | KIF20A |
| MCTS1 | SLC25A32 |
| PSRC1 | UTP11L |
| PSMA7 | SLC16A1 |
| ANLN | MRPL15 |
| TUBA1B | KIF4A |
| MAD2L2 | LRRC42 |
| GPI | PGK1 |
| TUBA1C | HK2 |
| MAP7D1 | AK3L1 |
| DDIT4 | CA9 |
| BNIP3 | |

#### IFN/AP

| |
|---|
| IFI6 |
| IFI35 |
| IFIT1 |
| IFIT3 |
| IFITM1 |
| IFITM3 |
| ISG15 |
| MX1 |
| OAS1 |
| STAT1 |
| STAT2 |
| TAP1 |
| B2M |
| CD74 |
| CIITA |
| HLA-B |
| HLA-DMA |
| HLA-DPA1 |
| HLA-DQA1 |
| HLA-DQB1 |
| HLA-DRA |
| HLA-DRB1 |
| HLA-DRB5 |
| HLA-E |
| HLA-F |
| NLRC5 |
